# Mapping Transcription Factor Networks By Comparing Tf Binding Locations To Tf Perturbation Responses

**DOI:** 10.1101/619676

**Authors:** Yiming Kang, Nikhil R. Patel, Christian Shively, Pamela Samantha Recio, Xuhua Chen, Bernd J. Wranik, Griffin Kim, Robi Mitra, R. Scott McIsaac, Michael R. Brent

**Author notes:** These authors contributed equally (Yiming Kang, Nikhil R. Patel). Corresponding author Michael Brent http://mblab.wustl.edu Campus Box 8510 Washington University Saint Louis, MO 63130.

## Abstract

**Background:** A transcription-factor (TF) network map indicates the direct, functional targets of each TF -- the genes it regulates by binding to their cis-regulatory DNA. Data on the genomic binding locations of each TF and the transcriptional responses to perturbations of its activity, such as overexpressing it, could support TF network mapping. Systematic data sets of both types exist for yeast and for human K562 and HEK293 cells.

**Results:** In previous data, most TF binding sites appear to be non-functional, so one cannot take the genes in whose promoters a TF binds as its direct, functional (DF) targets. Taking the genes that are both bound by a TF and responsive to a perturbation of it as its DF targets (*intersection algorithm*) is also not safe, as we show by deriving a new lower bound on the expected false discovery rate of the intersection algorithm. When there are many non-functional binding sites and many indirect targets, non-functional sites are expected to occur in the cis-regulatory DNA of indirect targets by chance. Dual threshold optimization, a new method for setting significance thresholds on binding and response data, improves the intersection algorithm, as does post-processing perturbation-response data with NetProphet 2.0. A comprehensive new data set measuring the transcriptional response shortly after inducing overexpression of a TF also helps, as does transposon calling cards, a new method for identifying TF binding locations.

**Conclusions:** The combination of dual threshold optimization and NetProphet greatly expands the high-confidence TF network map in both yeast and human. In yeast, measuring the response shortly after inducing TF overexpression and measuring binding locations by using transposon calling cards improve the network synergistically.

## BACKGROUND

Mapping out the circuitry by which cells regulate gene expression is a fundamental goal of systems biology. Such maps would facilitate a broad spectrum of research programs, much as maps of intermediary metabolism and genome sequences have done. Transcriptional regulation has multiple layers and component types, including sensors and signal transduction cascades involving kinases, phosphatases, and other enzymes. The bottom layer of transcriptional regulation, which acts directly at the genome, features sequence-specific DNA binding proteins known as transcription factors (TFs). Signaling cascades often change the activity levels of specific TFs -- the extent to which they exert their regulatory potential on their target genes -- via mechanisms that affect TFs’ abundance, localization, non-covalent interactions, or covalent modifications. To map and model transcriptional regulation as a whole, we must know which genes each TF regulates, or has the potential to regulate when activated.

A map of an organism’s TF network would have powerful applications. It could be used to infer the effects of specific signals, drugs, or environments on the activity levels of TFs by analyzing their effects on gene expression [1–4]. It could be used to predict the significance of naturally occurring genome variants in TFs or TF binding sites (TFBS). And, it could be used to design genome edits in TFs or TFBS to achieve a desired transcriptional state or behavior [5–7]. Crucial to all of these applications is the distinction between the direct functional targets of a TF -- the genes it regulates because it binds to their cis-regulatory DNA -- and its indirect targets, which are regulated via intermediary proteins. For example, a mutation inactivating a binding site for a TF in the cis-regulatory DNA of one of its direct targets will generally modulate or eliminate the relationship between the TF and its direct target. However, a mutation in a non-functional binding site which happens to lie in the cis-regulatory DNA of an indirect target will not affect the relationship between the TF and its indirect target.

In this paper, we analyze previously published and newly described genome-wide data sets, using both standard and novel analytic techniques, to reveal the current state of the art in identifying the direct, functional targets of a TF (Table 1). The data sets we focus on are those that aim to determine the binding locations of TFs and those that attempt to measure the transcriptional response to perturbations of TF activity, such as over expressing the TF or deleting the gene that encodes it. The binding location data derive from either chromatin immunoprecipitation (ChIP) or transposon calling cards, while the perturbation data include expression profiles after deletion of TF-encoding genes, short-term induction of TF expression, TF knockdowns via small-interfering RNA (siRNA) or small-hairpin RNA (shRNA), or CRISPR interference (CRISPRi). Yeast data sets are more complete than those of any other eukaryote and yeast has a simpler genome with more localized regulatory DNA, so we start by focusing on yeast. In addition to evaluating data sets and experimental and analytic methods, we construct a preliminary map of the yeast TF network by integrating the best available binding and perturbation response data sets. For model invertebrates, there are large data sets on TF binding location [8, 9], but there are currently no comparable data sets on the responses to perturbations of TF activity. We analyze large data sets on human cell lines from the ENCODE consortium [10, 11] and produce a preliminary, partial map of the TF network of human K562 cells. We also analyze a large data set on Zinc Finger TFs in human HEK293 cells [12].

**Table 1.** Data resources.

| Data type | Technology | Species | Proteins targeted | Targeted TFs analyzed | Genome assembly | Strain/Cell line | Publications |
| --- | --- | --- | --- | --- | --- | --- | --- |
| Binding location | ChIP-chip | *S. cerevisiae* | 203 | 155 | N/A | W303 | Harbison, 2004 |
|  | ChIP-chip | *S. cerevisiae* | 200 | 36 | N/A | S288C | Venters, 2011 |
|  | ChIP-exo | *S. cerevisiae* | 12 | 12 | R55, R64 (SGD) | S288C | Rhee, 2011; Rossi, 2018a; Rossi, 2018b; Bergenholm, 2018 |
|  | Transposon calling cards | *S. cerevisiae* | 15 | 15 | R61 (SGD) | S288C | Wang, 2012; Supplemental file S10 |
|  | ChIP-seq | *H. sapiens* | 261 | 261 | GRCh38 | K562 | Davis, 2018 (ENCODE) |
|  | ChIP-seq | *H. sapiens* | 131 | 131 | GRCh37 | HEK293 | Schmitges, 2016 |
| Perturbation response | TFKO | *S. cerevisiae* | 1,484 | 164 | N/A | S288C | Kemmeren, 2014 |
|  | ZEV TF induction | *S. cerevisiae* | 201 | 139 | N/A | S288C | Hackett, 2019 |
|  | TFKD (shRNA, siRNA) | *H. sapiens* | 261 | 261 | GRCh38 | K562 | Davis, 2018 (ENCODE) |
|  | CRISPRi | *H. sapiens* | 96 | 96 | GRCh38 | K562 | Davis, 2018 (ENCODE) |
|  | TF induction | *H. sapiens* | 80 | 80 | GRCh37 | HEK293 | Schmitges, 2016 |

Throughout most of this paper, we take the data sets at face value, assuming that they accurately report the molecular events they are designed to detect. An alternative explanation for some of our observations is that some data sets do not accurately report the events they are designed to detect. We do not take any position on that possibility nor do we mean to imply any judgment about it.

We consider several approaches to identifying the direct functional (DF) targets of a TF from binding location and perturbation-response data. The simplest is to take the genes in whose regulatory DNA a TF binds as its DF targets. We show that this is unsatisfactory as most binding sites for most TFs appear to be non-functional, according to existing data. We then consider taking the genes that are both bound by a TF and responsive to a perturbation of it as its DF targets, an approach we refer to as the intersection algorithm. We show, by means of a newly derived lower bound on the expected false discovery rate of the intersection algorithm, that it is unsatisfactory when applied to previously published data sets. Next, we introduce dual threshold optimization, a simple, new method for setting significance thresholds on binding and response data that improves the performance of the intersection algorithm. We then show, using a large, new perturbation-response data set, that measuring the response a short time after inducing the perturbation gives better results than measuring the steady-state response in a TF deletion strain. Next, we demonstrate that post-processing gene expression data through NetProphet 2.0 [13], a network inference algorithm, results in better agreement with binding data than using raw perturbation-response. Finally, we show that transposon calling cards, a recently developed method for identifying TF binding locations, improves the performance of the intersection algorithm greatly, especially when it is combined with rapid measurement of perturbation responses.

## RESULTS

### Simple comparison of yeast ChIP-chip to expression profiles of TF deletion strains yields few high-confidence regulatory relationships

#### Comprehensive binding and perturbation response data sets are available for yeast TFs

In 2004, Harbison et al. assayed the binding locations of essentially all yeast TFs by using ChIP-chip [14]. They identified many binding sites, but it was not possible at that time to estimate how many of them were functional, in the sense that the binding caused the TF to regulate the downstream gene. In 2007, Hu et al. published a data set in which all non-essential yeast TFs were deleted and the resulting deletion strains were subjected to expression profiling [15]. This made it possible for the first time to estimate the fraction of binding events that are functional, and Hu et al. remarked on how surprisingly small that fraction is -- about 3-5% in their data. In 2014, Kemmeren et al. published a second such data set, which benefited from newer technology and the hindsight afforded by the earlier study [16]. In the remainder of this paper, we focus on the Kemmeren TF knockout (TFKO) data because it demonstrates somewhat better agreement with the Harbison ChIP data, on average. We consider 183 yeast TFs with DNA binding domains and 5,887 genes that are labeled “verified” or “uncharacterized” in Saccharomyces Genome Database (SGD), omitting those that are labeled “dubious ORFS”.

#### Most bound genes in the Harbison ChIP data are not responsive in the TFKO data

We began this analysis by calculating the response rate of bound genes, for each TF -- the fraction of bound genes that are differentially expressed in the TFKO strain, relative to the wild type (WT). The spotted microarrays used by Harbison et al. in their ChIP-chip study contained one probe for each promoter, so their analysis yielded a simple P-value for whether each promoter is bound, with no further localization information. We eliminated from further consideration the 16 TFs that were not called as bound to any promoter. For the TFKO data, we used the authors’ statistical analysis and considered a gene to be differentially expressed in the TFKO strain, relative to the wild type strain, if its p-value (adjusted for multiple comparisons) was < 0.05. We eliminated from further consideration any TF whose knockout resulted in no significant changes as well as the 32 TFs whose microarray-reported expression level in the strain lacking the TF was more than one half its reported level in the WT. This can happen when the wild-type expression level of the TF is near or below the detection limit of the microarray.

Fig. 1A shows a histogram of the results. The median response rate for bound genes was 18%. The mode was 0% -- 25 of the 97 TFs (26%) had both bound targets and responsive targets, but none of the bound targets were responsive. Only 17 TFs (18%) had a response rate above 50%. This is without requiring a minimum fold change, to filter for biologically significant responses. With a minimum fold change of 1.5, the number of TFs for which more than half the bound genes are responsive drops to 4 (4%; Fig. S1A). Tightening the significance threshold for binding to P<10^−5^ with no minimum fold change for response increases the median response rate to ~28%, but at the cost of reducing the total number of bound and responsive genes, summed over all TFs, to 297 (Fig. S1A-C). Thus, these data do not support the notion that most binding is functional. The low response rate of bound genes cannot be explained by saying that the TFs are inactive in the conditions tested, since the median number of genes that respond with p<0.05 is 321 (13 with fold-change >1.5). A lot of genes respond, they’re just not the same ones that are bound.

**Figure 1.**
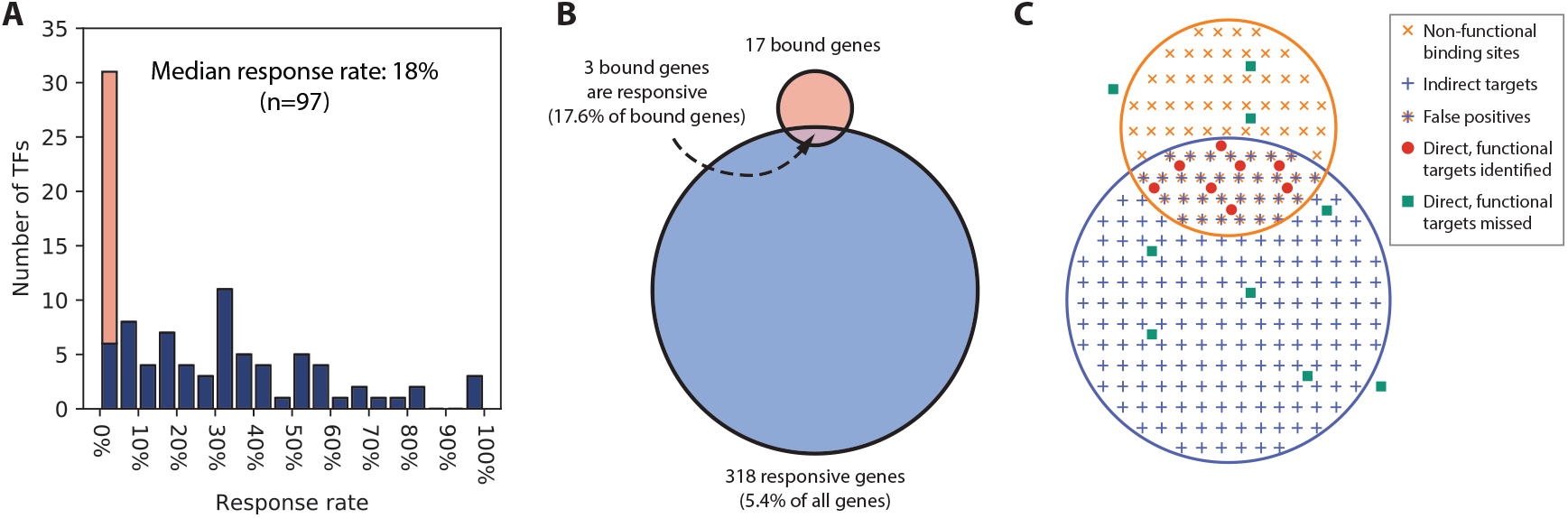
(A) Distribution of the response rates of TFs (fraction of bound genes that respond to TF perturbation) in the Harbison binding and Kemmeren TFKO data sets. Stacked orange bar indicates the number of TFs with response rates of exactly 0. Binding threshold is p<0.001 and response threshold is p<0.05, as recommended in the original publications, with no minimum fold change. (B) Median numbers of bound genes (17), perturbation-responsive genes (318), and intersection size (3), when comparing the ChIP-chip data to the TFKO perturbation-response data. Thresholds are as in panel A. (C) An illustration of the idea that non-functional binding sites will sometimes occur in the promoters of responsive, indirect targets by random chance.

#### Many genes that are both bound and responsive in previously published data are probably not DF targets

Given that available data suggest most binding sites are non-functional, a logical procedure for finding the DF targets is to choose those that are. In other words, to take the intersection of the genes bound by each TF with the genes that respond to a perturbation of that TF. It is important keep in mind, however, that most responsive genes are not bound. Comparing the ChIP data with the TFKO data, the median fraction of responsive genes that are bound is 1% (Fig. 1B). Thus, most of the responsive genes are indirect targets. Furthermore, it is reasonable to assume that the distribution of indirect targets among all genes is independent of the distribution of non-functional binding sites (Fig. 1C). Or at least that non-functional binding sites do not systematically avoid the promoters of indirect targets. This suggests that some of the indirect targets also have non-functional binding sites. These genes would be false positives of the intersection algorithm -- genes that are bound and responsive, but are not responsive *because* they are bound.

In Box 1, we derive a lower bound on the expected false discovery rate (FDR) of the intersection procedure, as a function of the number of bound genes, |B|, the number of responsive genes, |R|, the number of bound and responsive genes, |B∩R|, and the total number of genes, |G|. The lower bound also depends on the sensitivity of the intersection procedure -- the fraction of direct functional targets that are in the intersection. The formula shows that, if a large fraction of bound genes are not responsive and a large fraction of responsive genes are not bound, the intersection procedure cannot have both high sensitivity and low false-discovery rate. For example, Fig. 2A shows the relationship between sensitivity and expected FDR for a fairly typical TF, Gln3, based on the Harbison ChIP data and the TFKO response data. A reasonable minimum accuracy criterion for a procedure aimed at finding the DF targets of a TF is that it have sensitivity >= 80% (it detects at least 80% of the DF targets) and an FDR <= 20%. However, that is not possible for Gln3, using these two data sets. Intuitively, this is because the fraction of Gln3-bound genes that are responsive to the Gln3 perturbation (53%) is only a little more than the fraction of all genes that are responsive to the Gln3 perturbation (43%; Fig. 2B). The 80-20 criterion is achievable for only 43 TFs. Fig. S2 shows the cumulative fraction of TFs that have an FDR bound below a given level, assuming 80% sensitivity, at various significance thresholds for binding and response.

**Figure 2.**
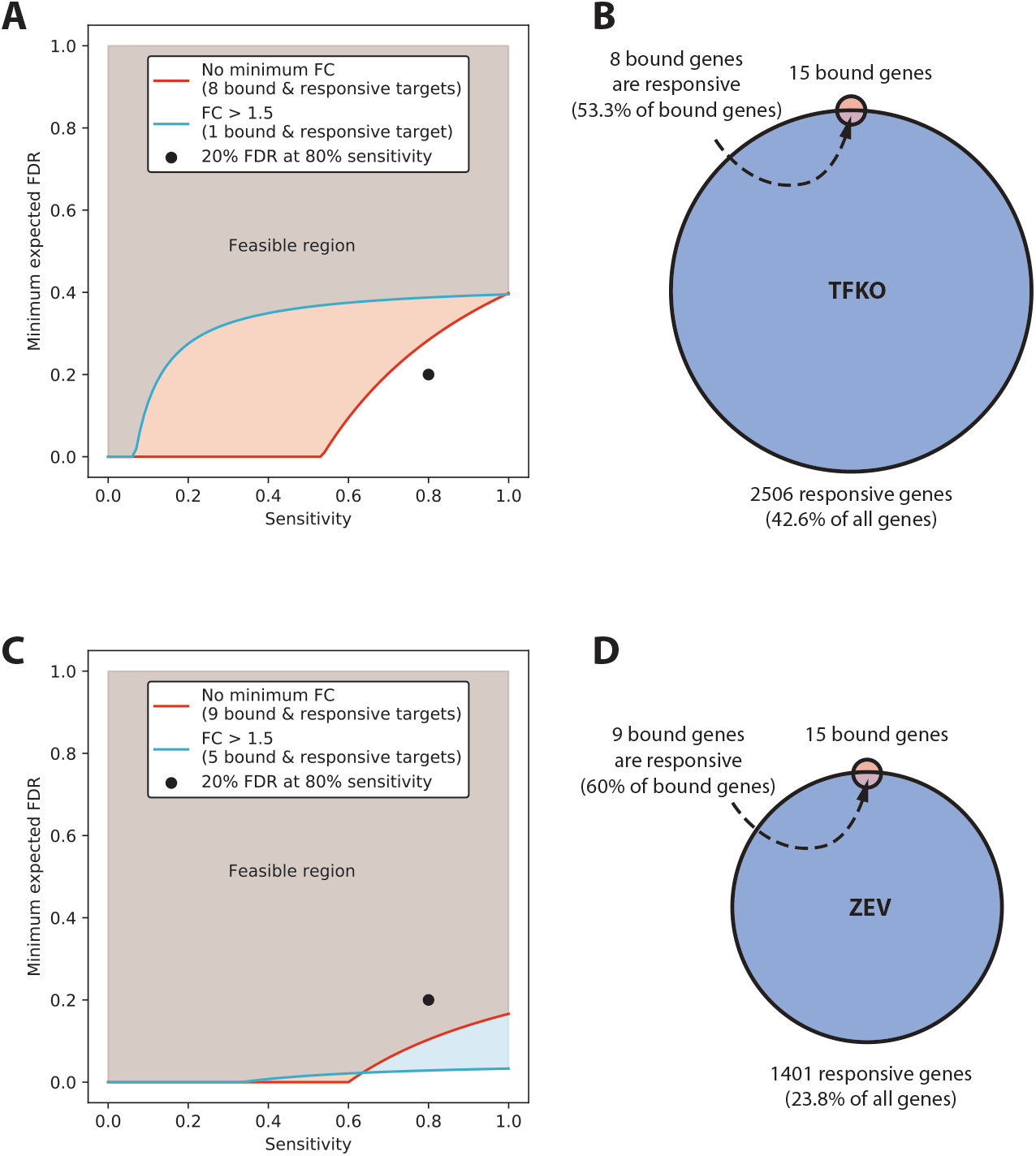
Analysis of Minimum expected false discovery rate (FDR). (A) Minimum expected FDR as a function of sensitivity for TF Gln3, with moderate and tight thresholds for responsiveness, when comparing ChIP to TFKO. 80% sensitivity with 20% FDR is not attainable at either threshold, when comparing ChIP to TFKO. (B) The bound set, responsive set, and intersection for Gln3, when comparing ChIP to TFKO. (C) Minimum expected FDR, as a function of sensitivity, with moderate and tight thresholds for responsiveness, when comparing ChIP to ZEV15. 80% sensitivity with 20% FDR is attainable at either threshold. (D) The bound set, responsive set, and intersection for Gln3, when comparing ChIP to ZEV15.

###### Box 1: Expected False Discovery Rate (FDR) of intersection algorithms

Intersection algorithms identify the direct functional targets of a TF as those whose promoters are bound by the TF in an assay such as ChIP-Seq and are responsive when the same TF is perturbed. A true direct functional (DF) target is responsive when the TF is perturbed *because it is bound by the TF*. The obvious alternative is that the binding site is non-functional and the gene is responsive because it is an indirect target of the TF. Another possible alternative is that the gene is a false positive of the binding or response assay.

We start by defining the following notation for any given TF:

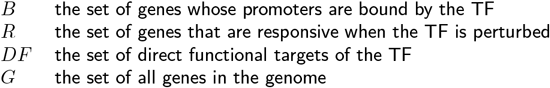

The analyses below are based on a hypothesis that is best understand by first thinking about the non-functional binding sites of a TF and its indirect targets as being distributed randomly and independently across the genes that are not DF targets. In notation:

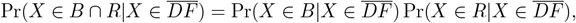

That is, having a non-functional binding site for a TF and being an indirect target of the TF are unrelated – an indirect target is no more or less likely to have a non-functional binding site than any other gene. However, our proof does not require equality, just the inequality

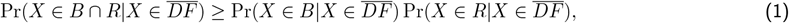

where *X* is a randomly chosen gene. That is, indirect targets and non-functional binding sites do not systematically avoid one another.

The sensitivity of the intersection algorithm is:

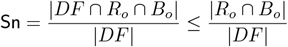

where the subscript *o* emphasizes that we are referring to the actual observed sets of bound and responsive genes from some particular experiment. Thus,

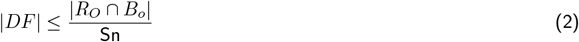

The expectation of the FDR, with respect to the random process that distributes non-functional binding sites and indirect targets, is

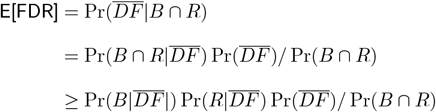

where 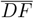 is the set complement of *DF* and the random variable *X* has been omitted for brevity.

We can estimate these probabilities by maximum likelihood from the observed bound and responsive sets, *B*_*o*_ and *R*_*o*_, as follows.

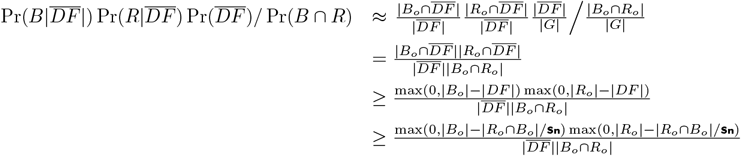

Based on these estimates

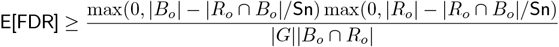

The FDR lower bound is only a lower bound and does not guarantee any maximum FDR for the intersection algorithm. In fact, of the 43 TFs that could possibly achieve the 80-20 criterion in the ChIP-TFKO comparison, only 27 have an intersection that is significantly larger than would be expected by chance (hypergeometric P<0.01, not adjusted for multiple testing). If we define “acceptable TF” to be one that could pass the 80-20 criterion and has a larger overlap between bound and responsive targets than would be expected for randomly selected gene sets, then there are 27 acceptable TFs with a total of 448 regulatory interactions involving 366 unique target genes. If we take this to be our network map, ~85% of TFs are not acceptable so they have no high confidence targets, while 94% of genes have no identifiable regulator. Clearly, using the simple intersection algorithm with just these two data sets is not producing anything like a complete network map.

### Comparing yeast ChIP-chip data to expression profiling shortly after TF induction enlarges the map

Recently, Hackett et al. released a data set in which the expression of nearly every yeast TF was induced from a very low level to a high level [17]. This was accomplished by expressing ZEV, an estradiol-activated artificial TF, and replacing the promoter of the gene to be induced with a ZEV-responsive promoter [18, 19]. (Some of the TFs were induced using an earlier iteration of the artificial TF called GEV [20], but we refer to the data set as ZEV for convenience.) Gene expression profiles were measured before induction and at 5, 10, 15, 20, 30, 45, and 90 minutes after inducing the expression of a natural yeast TF with estradiol. We reasoned that genes that respond rapidly might be enriched for direct targets of the induced TF, since there would be limited time for intermediary proteins to be transcribed and translated. If the responders were enriched for direct targets, the number of acceptable TFs might increase, expanding the network map. In general, the expression profiles taken 15 minutes after TF induction (ZEV15) were most enriched for bound genes, so we focus on the 15-minute time point for the remainder of the paper. Specifically, among the 94 TFs available in Harbison ChIP and all ZEV time points, ZEV data at 15 minutes yielded the maximal number of acceptable TFs when compared to ChIP data (Fig. S3). We consider a gene to be responsive if its shrunken log fold change estimate, relative to time 0, was non-zero for details of the shrinkage analysis). A detailed description of these strains and expression profiling experiments can be found in ref. [17]

The TF Gln3, which could not achieve 80% sensitivity with 20% expected FDR in the ChIP-TFKO comparison (Fig. 2A), can in the ChIP-ZEV15 comparison (Fig. 2C). The reason is that the number of responsive genes has decreased from 43% of all genes to 24%, at the same time that the response rate of bound genes *increased* from 53% to 60% (Fig. 2B, D). Across all TFs, the ChIP-ZEV15 comparison identified 37 acceptable TFs, 23 of which had not been identified in the ChIP-TFKO comparison (Fig.3A). Together, the ZEV15-ChIP and TFKO-ChIP comparisons yielded 50 acceptable TFs with a total of 930 regulatory interactions involving 722 unique target genes. This network map is significantly expanded, but it is still the case that >72% of TFs are not acceptable and hence have no targets, while >87% of genes have no regulators.

**Figure 3.**
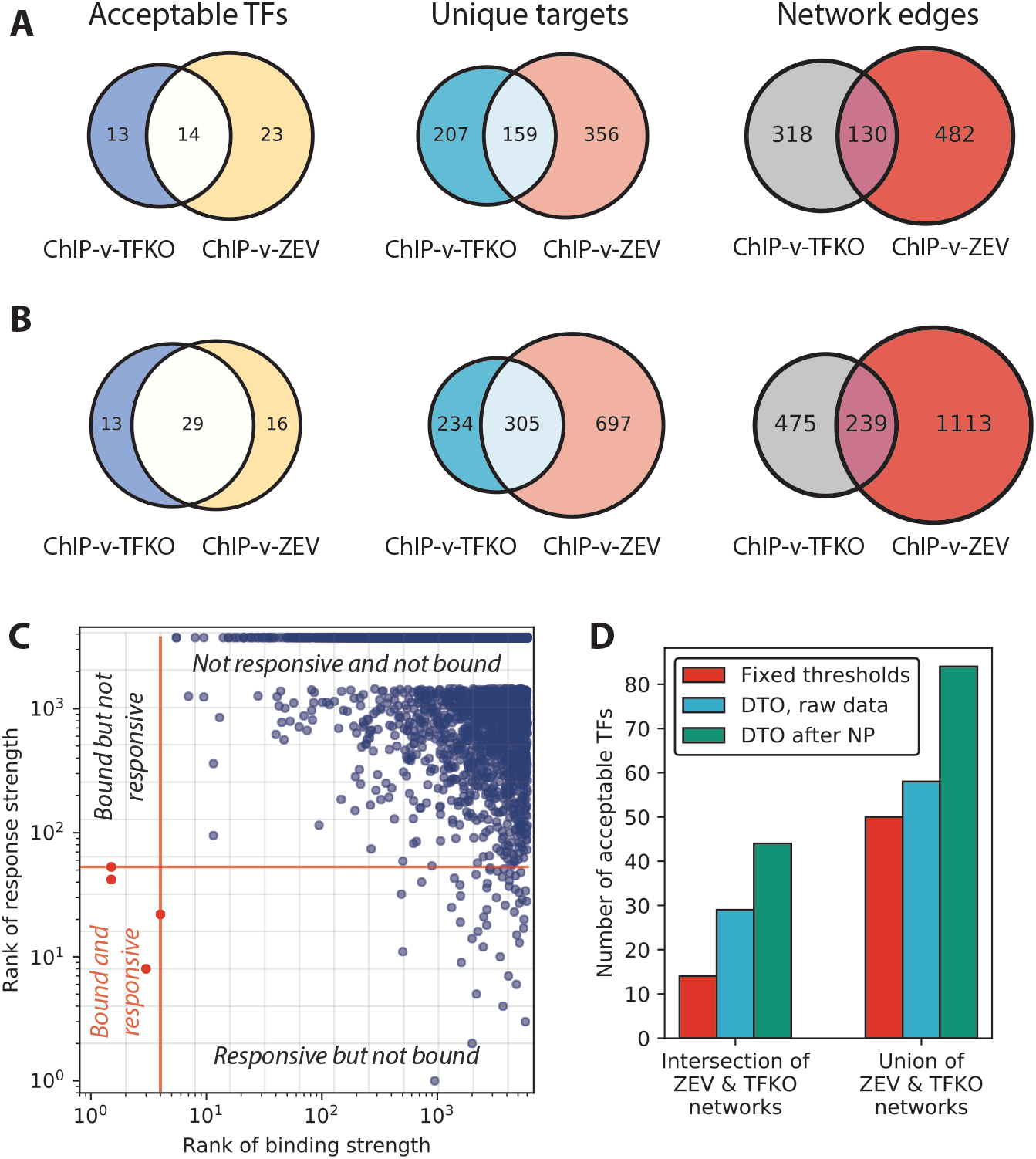
(A) Numbers of acceptable TFs, unique target genes, and network edges, when comparing Harbison ChIP data to TFKO or ZEV15 response data. “Unique Targets” are genes that are in the bound-responsive intersection of an at least one acceptable TF and thus are plausible direct functional targets. Edges connect acceptable TFs to the genes in their bound-responsive intersection. These genes are not guaranteed to be direct functional targets. The ZEV15 response data yields more acceptable TFs, more regulated genes, and more regulatory edges. (B) Numbers of acceptable TFs and unique target genes for comparison of Harbison ChIP binding data to TFKO or ZEV15 response data, after dual threshold optimization. The requirement that the overlap between the bound and responsive targets be significantly greater than chance at P<0.01 was obtained by comparing the nominal hypergeometric P-value for the overlap to a null distribution obtained by running dual threshold optimization on 1,000 randomly permuted binding and response data sets. ZEV yields more acceptable TFs, regulated genes, and regulatory interactions than TFKO. (C) Illustration of DTO algorithm. Each dot represents one gene. Red lines indicate the chosen (optimal) thresholds for binding (vertical red line) and regulation (horizontal red line). The lower left quadrant, relative to the red lines, contains the bound and responsive genes, which are presumed to be direct functional targets (red dots). The lower right quadrant contains genes that are judged to be responsive but not bound and the upper left quadrant contains genes that are judged to be bound but not responsive (in this case there are none). Gray lines indicate some of the other possible thresholds on binding or response and locations where the gray lines cross are possible combinations of binding and response thresholds, each of which is evaluated by the DTO algorithm. (D) Comparison of TFKO and ZEV15 networks derived from fixed thresholds, DTO on raw gene expression, and DTO on gene expression data processed by NetProphet 2.0. The use of DTO on the raw expression data (blue bars) increases the size of the networks, whether you focus on the intersection of the TFKO and ZEV networks (left bar grouping), or the union (right bar grouping). Post processing with NetProphet 2.0 (green bars) increases the number of acceptable TFs.

### Dual threshold optimization expands the TF Network map

A possible limitation of the previous analyses is that they rely on statistical significance thresholds to determine which genes are bound and which are responsive. The statistics are calculated separately for the binding and response data sets and statistical significance threshold are, by their nature, arbitrary. Furthermore, statistically significant levels of binding or perturbation response might not be biologically significant. For example, a TF may bind a site consistently in the ChIP data even though the fractional occupancy of the site is too low to detectably affect transcription. To address these problems, we developed dual threshold optimization (DTO), a method that sets the binding and response thresholds by considering both data sets together. DTO chooses, for each TF, the pair of (binding, response) thresholds that minimizes the probability that the overlap between the bound and responsive sets results from random gene selection (Fig. 3C).

For this analysis, we ranked all genes by their absolute log fold change in the ZEV15 data and, separately by their negative log P-value in the Harbison ChIP data. We could have used the underlying ChIP signal rather than its P-value, but in this case the P-value was more convenient (see below). The genes with the strongest evidence for binding or responsiveness were ranked at the top of the lists. We then chose the pair of (binding, response) rank thresholds so as to minimize the probability of the overlap between bound and responsive, under a null hypothesis of random selection of gene sets of the sizes determined by the thresholds (hypergeometric distribution). The only constraint on the thresholds chosen was that the P-value for the ChIP data could not exceed 0.1.

To test the significance of the overlap at the chosen thresholds, we needed a null distribution for the results of running DTO on unrelated binding and response rankings. The null distribution for randomly chosen, fixed-size sets does not apply because DTO chooses the bound and responsive set sizes specifically to minimize probability under the fixed-size null. To obtain the correct null, we randomly permuted the assignment of binding and response signals to genes 1000 time for each pair of binding and response data and ran DTO on each random permutation (see Supplemental Methods for details).

After DTO, we applied the same acceptability criteria as before -- the bound and responsive overlap must be significant (P<0.01, permutation-based) and 20% FDR at 80% sensitivity must be achievable. DTO expanded the network map again (Fig. 3B). Combining the results of both response data sets, it yielded 58 acceptable TFs with a total of 1,829 regulatory interactions involving 1,236 unique target genes. Interestingly, the number of TFs that are acceptable in both data sets now exceeds the number that are acceptable in either of the data sets alone. In this map, ~32% of TFs have at least one target and ~21% of genes have at least one regulator. The maps based on DTO of TFKO and ZEV15 data are provided as Supplemental Files S1 and S2.

### Processing yeast gene expression data through NetProphet 2 further expands the map

NetProphet 2.0 [13] is a network inference method that combines analysis of gene expression data, including expression data from perturbation-response experiments, with information gleaned from genome sequence. It assigns a score to each possible TF-target interaction and ranks all possible interactions according this score. A major component of the NetProphet score is the degree to which the target gene responds to direct perturbation of the TF. However, it also considers the degree to which the mRNA level of the TF is predictive of the mRNA level of the potential target, across many different perturbations. This analysis is not limited to single-TF perturbations -- it can also use perturbations of other genes, drugs, or growth conditions. As a result, it can make predictions about the targets of TFs that have not been individually perturbed. NetProphet also makes use of two other ideas: (1) that co-regulated genes tend to have similar sequence motifs in their promoters, and (2) that DNA binding domains with similar amino acid sequences tend to bind similar motifs. NetProphet 2.0 combines all these factors, but is primarily driven by gene expression data. It does not use ChIP or other experimental data on TF binding location.

We built separate NetProphet networks using the TFKO and ZEV data (Methods). For TFKO, we input 3 wild-type expression profiles and the complete set of 1,484 expression profiles from strains lacking one gene -- some of the deleted genes encode TFs, but others encode other putative regulatory proteins, such as kinases and phosphatases. For ZEV, we used 590 expression profiles from 15 minutes, 45 minutes, or 90 minutes post-induction. We then ranked the potential targets of each TF by their NetProphet scores and ran dual threshold optimization, treating the NetProphet score as we did the response strength.

Combining the results from NetProphet applied to TFKO and NetProphet applied to ZEV, dual threshold optimization resulted in 84 acceptable TFs (Fig. 3D) with a total of 2,151 regulatory interactions (Fig. S4B) involving 1,326 unique target genes (Fig. S4A). The number of TFs that are acceptable in both data sets, 44, is now much larger than the number that are acceptable in either data set alone (TFKO:22, ZEV:18). The total number of edges, combining both TFKO and ZEV data, is 2,151, of which 400 are supported by both data sets.

Supplemental Files S3 and S4 contain the complete set of regulatory edges for each acceptable TF in each comparison, along with their NetProphet score rank (among all possible interactions), their Harbison ChIP P-value, and the probability of the TF’s bound-responsive intersection under a hypergeometric null model. This may be the best network that can be obtained by using the comprehensive yeast ChIP-chip data. Network inference methods that do not consider binding data, such as NetProphet 2.0, may produce better networks on their own, but if support from binding data is required, this is the best we can do with these data sets. In this network, ~46% of TFs have at least one target and ~37% of genes have at least one regulator. Running NetProphet on gene expression data and feeding the result into dual threshold optimization has enlarged the map, but it is still smaller than what is generally expected for the complete yeast TF network. To improve on it further, we need binding data that is more focused on functional binding or simply more accurate. We will consider newer yeast binding data, produced by using new methods, after discussing ChIP-seq and perturbation response data on human cells lines.

### Without processing by NetProphet 2.0, data on human cell lines yields a few acceptable TFs

The ENCODE project [10] has produced a wealth of data on human cell lines, which currently includes 743 TF ChIP-Seq experiments and 391 RNA-Seq experiments following knockdown of a TF by siRNA or shRNA (TFKD) or repression of a TF by CRISPR interference (CRISPRi) [21]. In this section, we refer to proteins as TFs if they were subjected to both ChIP-Seq and perturbation-response experiments and are listed as TFs in the ENCODE database; the question of which proteins are sequence-specific DNA binding proteins that regulate transcription rates is considered further in the Discussion. In K562 cells, 42 TFs have both ChIP-Seq and TFKD data and 45 TFs have both ChIP-Seq and CRISPRi data. In HepG2 cells 16 TFs have both binding and TFKD data. We focus our investigation on K562 data, as it is by far the biggest relevant data set.

We considered two ways of assigning ChIP-Seq peaks to the genes they potentially regulate. The first is the traditional approach of choosing an interval around the transcription start site (TSS) -- we used 10 kb upstream to 2 kb downstream. The second is to take a small proximal promoter region (TSS −500 bp to +500 bp) along with enhancer regions that have been identified and assigned to the target gene in the GeneHancer database [22]. GeneHancer uses a variety of data types including predicted and ChIP-based TF binding sites, enhancer RNAs, histone marks, chromosome conformation, and cis-EQTLs. We used only the ‘elite’ enhancers and ‘elite’ associations, each of which are supported by at least two sources of evidence. In order to be comprehensive, we used all elite enhancers and enhancer-gene associations, regardless of the cell lines or tissue types in which the evidence was obtained. However, 91% of the ‘elite’ enhancers were supported by evidence from ENCODE, much of which comes from the same K562 cell line used for the binding and perturbation-response studies. The enhancer-based approach generally gave 1 or 2 more acceptable TFs than the fixed interval, so we used that in subsequent analyses.

Unlike the yeast array-based data, the human sequencing-based data tended to yield many more bound than responsive genes (Fig. 4A, B). Among the TFs that had at least one bound and one responsive gene, 7 (TFKD) and 7 (CRISPRi) had no genes that were both bound and responsive. The median response rate for bound genes was <0.5%. In a fixed-threshold comparison to K562 ChIP-Seq data with adjusted P < 0.05, TFKD and CRISPRi produced 5 acceptable TFs each. We then ran dual threshold optimization limiting the bound and responsive gene sets to have P<=0.1; such limits are necessary because DTO occasionally chooses implausible thresholds, such as counting all genes as responsive. DTO increased the number of acceptable TFs slightly, to 6 and 6. In these data sets, all TFs that failed the 80-20 FDR criterion also failed the overlap P<0.01 criterion, so the results would be the same without the FDR criterion. However, all TFs would have failed if we had required 80% sensitivity and FDR <= 10%. Of the total number of TFs with both binding and response data, TFKD yielded 14% acceptable TFs (6/43) and CRISPRi yielded 13% (6/45).

**Figure 4.**
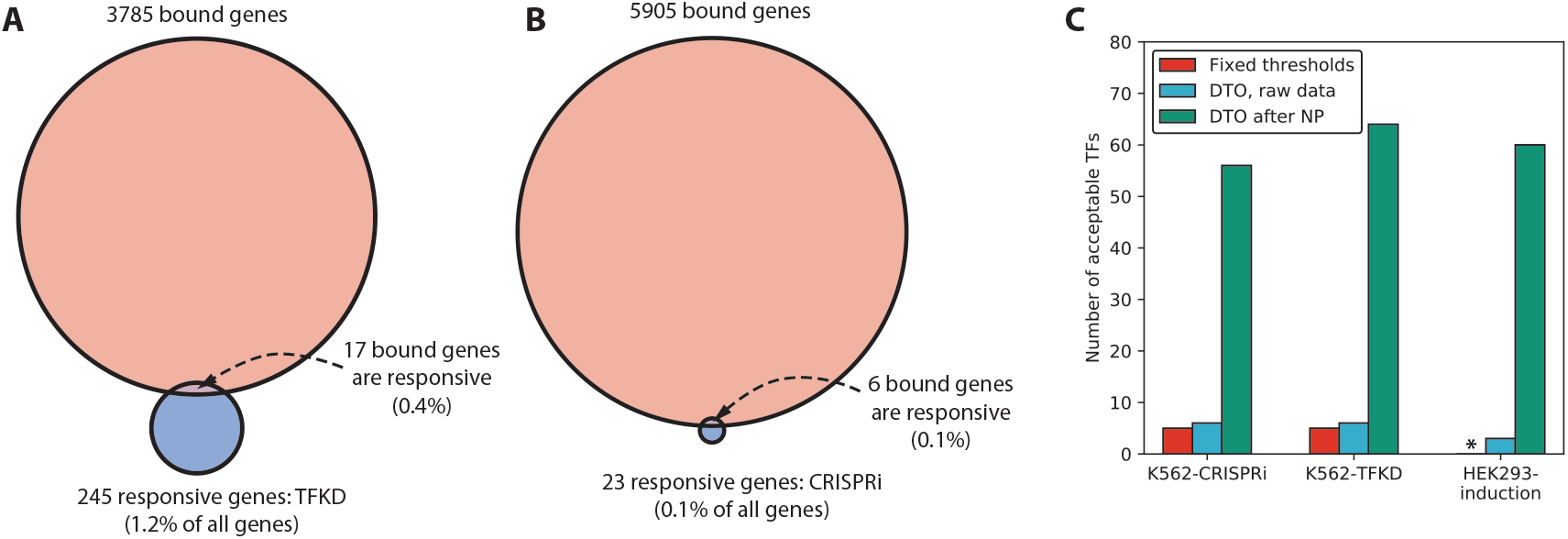
(A) Medians of number of bound genes, number of perturbation-responsive genes, and number genes that are both bound and responsive, when comparing ENCODE K562 ChIP-Seq data to ENCODE TFKD data. Excludes TFs with either no bound genes or no responsive genes. Binding threshold is p<0.05 and response threshold is p<0.05 with no minimum fold change. (B) Comparison of ENCODE K562 ChIP-Seq data and ENCODE CRISPRi data, as in Panel A. (C) Comparison of human networks derived from fixed thresholds, dual threshold optimization (DTO) on raw gene expression, and DTO on gene expression data processed by NetProphet 2.0. No fixed threshold analysis for HEK293 is available for the lack of response p-value.

We also analyzed a data set on 88 human GFP-tagged C2H2 Zinc Finger TFs with matched Chip-Seq data and response-to-overexpression data in HEK293 cells [12]. ChIP-Seq was carried out using an antibody against GFP and RNA-Seq was carried out 24 hours after overexpressing the TF from a tetracycline-inducible plasmid. For the majority of TFs there was only a single replicate of the RNA-Seq experiment, which prevents the calculation statistical significance by traditional methods. However, we were able to carry out DTO using the absolute log fold-change in the single replicate (relative to the median expression in all perturbations) as the measure of response strength. Seven of the 88 TFs were acceptable in this analysis, but DTO chooses all genes as responsive for five of the seven. When we limited the total number of responsive genes to 300,000, or 3409 per TF on average, three TFs were acceptable and none had more than 1,000 responsive genes.

### Processing human data through NetProphet 2.0 greatly increases the number of acceptable TFs

We ran NetProphet 2.0 on both the K562 data (TFKD and CRISPRi) and the HEK293 data described above (see Supplemental Methods for details). We then ran DTO limiting the total set of responsive genes to those with the top 500,000 (K562) or 300,000 (HEK293) NetProphet scores. NP can infer targets for TFs that have not been directly perturbed by exploiting correlation between the expression of the TF and its targets (among other factors), so we were able to calculate NP scores for 262 ChIPed TFs (K562) or 103 ChIPed TFs (HEK293). This greatly increased the number of acceptable TFs to 64 for K562 TFKD (24%) and 56 for K562 CRISPRi (21%; Fig. 4C). For HEK293, DTO on NetProphet scores increased the number of acceptable TFs to 60 of the 103 that were chipped (58%).

### More recent yeast ChIP data does not yield as many acceptable TFs as the Harbison data

The ChIP data published by Harbison et al. in 2004 is still the only data set of binding locations of most or all yeast TFs. However, Venters et al. [23] carried out ChIP-chip on a set of proteins they termed chromatin factors, 25 of which were also chipped by Harbison and perturbed by TFKO and ZEV. We carried out dual threshold optimization on these 25 common TFs, comparing the two ChIP binding data sets to the TFKO and ZEV perturbation data sets, with and without post-processing by NetProphet 2.0. The older Harbison ChIP data produced more acceptable TFs than the Venters data, when compared to either TFKO or ZEV, either with or without NetProphet (Fig. 5A). Thus, the age of the Harbison ChIP data does not seem to be a significant limitation.

**Figure 5.**
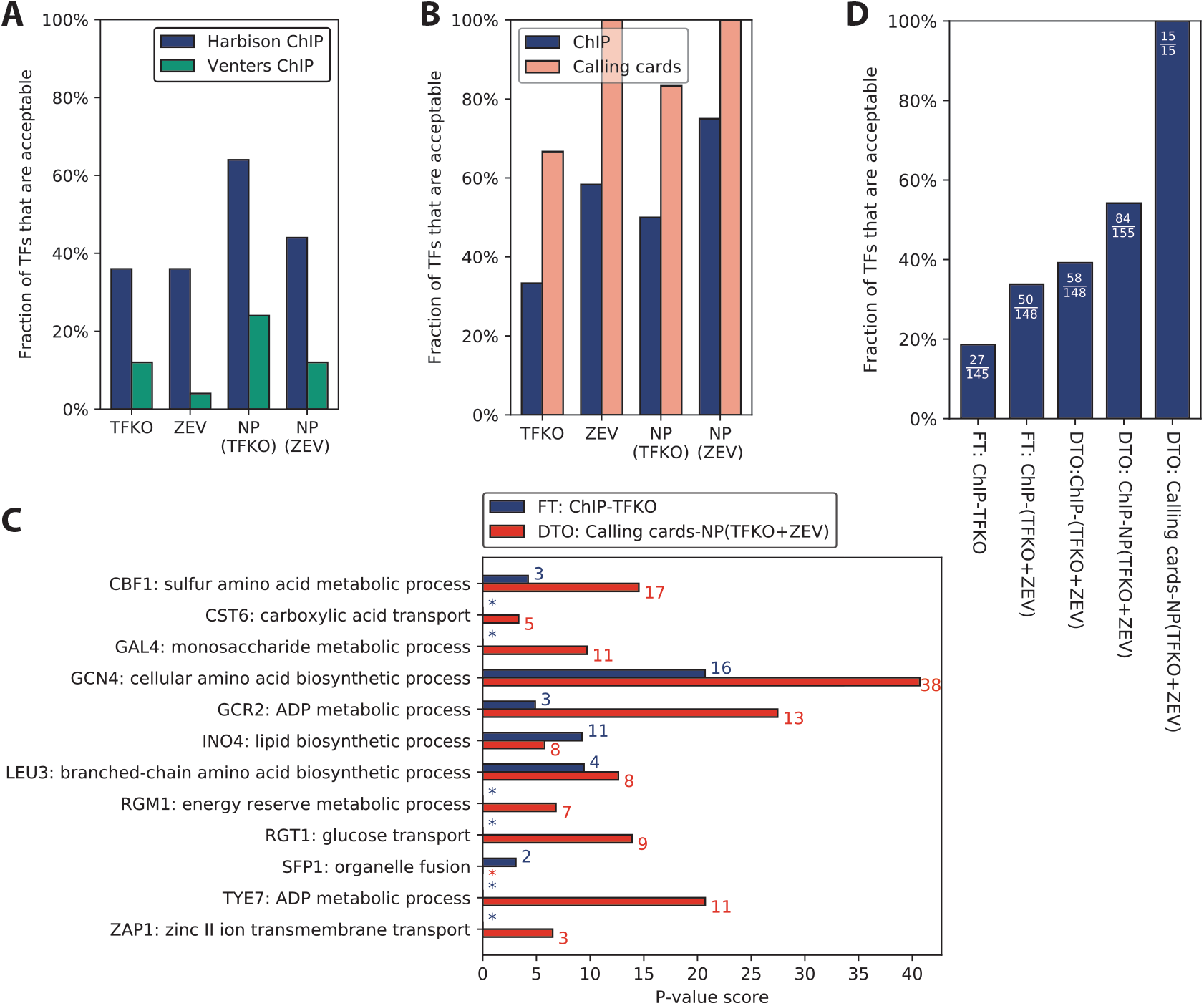
(A) Number of acceptable TFs, comparing the Harbison ChIP and Venters ChIP data on the same 25 TFs. Regardless of the perturbation data set or the processing by NetProphet 2.0, the Harbison ChIP data always yields more acceptable TFs. (B) Among the 12 TFs for which we have data in Harbison ChIP, calling cards, TFKO, and ZEV, the percentage that are acceptable. Regardless of the perturbation data set or processing by NetProphet 2.0, calling cards always yields more acceptable TFs. Holding all other factors constant, ZEV always yields more acceptable TFs than TFKO. NetProphet postprocessing always yields more acceptable TFs than raw differential expression for TFKO; for ZEV, the raw data already achieve 100% acceptable. (C) For each of the 12 TFs for which we have data in Harbison ChIP, calling cards, TFKO, and ZEV, the gene ontology (GO) term that is most strongly enriched in its targets. Targets are determined either by simple intersection of the bound and responsive genes in Harbison ChIP and TFKO data, using fixed thresholds (blue) or by dual threshold optimization on calling cards data and output from NetProphet 2.0 run on the TFKO and ZEV expression data (red). The colored numbers indicate the number of target genes annotated to the most significant GO term. Asterisk indicates no GO enrichment with P<0.01. (D) Among all TFs for which the indicated analyses can be carried out, the percentage that are acceptable in either TFKO or ZEV expression data or both. The fraction shows the number of acceptable TFs over the total number of TFs that could be analyzed. FT: Fixed threshold. DTO: Dual threshold optimization.

### ChIP-exo yields more acceptable TFs than traditional ChIP

We also ran NetProphet and DTO on a small set of TFs for we could obtain binding data from ChIP-exo, a variant of the ChIP method in which the affinity-purified chromatin is digest by DNase, leaving a much smaller piece that is partially protected by protein. Seven TFs had data in ChIP-exo, Harbison, TFKO, and ZEV, enabling all-way comparisons. Regardless of the perturbation-response data set, ChIP-exo always had more acceptable TFs than ChIP-chip (Fig. S5). However, five TFs had ChIP-exo data in four different growth conditions. We used the nitrogen-limited chemostat data as it gave the best results, however this may overestimate the agreement that would be found in a more typical scenario where ChIP-exo is performed in only one condition. After processing either the ZEV or TFKO perturbation-response data through NetProphet 2.0, all seven TFs were acceptable (Fig. S5). Thus, while the numbers of TFs are still small, this analysis suggests that ChIP-exo may yield better agreement with perturbation-response data than traditional ChIP.

### Transposon calling cards yields more acceptable TFs than traditional ChIP

Transposon calling cards is a method of determining TF binding locations by tethering a transposase to a TF, recovering the inserted transposons with their flanking sequences, and counting the insertions in a given genomic region. It does not require crosslinking, sonication, or affinity purification (see refs [24–26] for details). Here, we analyze both previously published [24] and new, previously unpublished calling cards data. Binding data from ChIP and Calling Cards were compared to perturbation-response data from TFKO and ZEV, using the 12 TFs present in all 4 data sets (Fig. 5B). In all comparisons, calling cards yielded substantially more acceptable TFs than ChIP. This is particularly impressive given that the calling cards experiments were carried out very different growth conditions from the ZEV experiments -- synthetic complete medium with galactose on agarose plates at room temperature versus rich medium with glucose in liquid culture at 30C. Figure 5B also shows that, holding all other factors constant, ZEV was always better than TFKO and post-processing by NetProphet was always beneficial. Lists of acceptable TFs and their bound and responsive targets for all calling cards analyses in Fig. 5B are provided as Supplemental Files S5-8.

Figure 5C shows the -log P-value of the most significant gene ontology (GO) term for the predicted targets of each TF we have calling cards data on, excluding terms that describe more than 300 or fewer than 3 genes. To highlight the progress reported here, results are shown for the best combination of experimental and analytic methods (DTO on calling cards data and NetProphet output after running on TFKO and ZEV 15, 45, and 90-minute samples) and for the simple intersection of bound and responsive genes using TFKO and ChIP-chip. For 10 of 12 TFs, the best combination of methods had a stronger GO term P-value, and the differences were large. For 2 of 12 (Ino4 and Sfp1), simple intersection had the stronger P-value, but the differences were smaller. The median -Log10 P-value for the best combination of methods was 11.2, while that of simple intersection was 1.5. The best combination of methods assigned the top GO term to 117 target genes, whereas simple intersection assigned the top term to only 41 genes. For most TFs, the most significant GO term had a clear relationship to the known function of the TF as described in the Saccharomyces Genome Database. This includes some cases where the term selected is an immediate parent of the most familiar term associated with the TF. For example, Gcr2 (Glycolysis Regulation 2) is known as a regulator of genes encoding glycolytic enzymes. Its most significant GO term is “ADP metabolic process”, annotating 13 predicted Gcr2 targets, but 12 of those targets are also annotated with “Glycolytic process”, a child (subcategory) of “ADP metabolic process”. This can be seen in Figure S6, which shows the top 5 GO terms for each TF.

Another way to look at the contributions of various methods is to plot the fraction of available TFs that are acceptable, combining TFKO and ZEV, using each combination of methods described here (Fig. 5D). Only 15 TFs are currently available for calling cards and either ZEV or TFKO (12 for both), but analyzing these with DTO and NetProphet results in a much larger fraction of TFs being acceptable. This includes TFs that are not thought to be active in the ZEV or TFKO growth conditions, such as Gal4, presumably because ZEV overexpression of Gal4 significantly exceeds the number of Gal80 molecules available to bind and inactivate it.

### The combination of ZEV and calling cards greatly increases response rates

We began this paper by observing that, using fixed threshold analysis of the TFKO and ChIP data, most binding appears to be non-functional. To revisit the question of functionality using ZEV and calling cards data, we plotted the fraction of bound genes that are responsive, as a function of binding strength rank. Figure 6A shows that, for the TF Leu3, the combination of calling cards and ZEV gives much higher response rates than any of the other three combinations -- ChIP-ZEV, calling cards-TFKO, or ChIP-TFKO -- regardless of binding strength. Nine out of the 10 mostly strongly bound and 48 out of 100 most strongly bound genes were responsive. To make the comparison between ZEV and TFKO fair, we fixed the number of Leu3-responsive genes in each data set to be the same. Thus, we labeled the 156 most strongly responsive genes in each data set as responsive. We chose 156 because it was the minimum of the numbers of genes that were significantly differentially expressed in the two data sets for Leu3. Although the number of responsive genes in each data set was the same, a larger fraction of the ZEV-responsive genes was bound, as compared to the TFKO-responsive genes. Figure 6B shows a similar plot of the average response rates at each binding threshold, across the 12 TFs for which we have all four combinations of data sets. Again, the combination of ZEV and calling cards gives higher response rates at all binding thresholds. On average, the response rate of the 10 most strongly bound genes is 56%. However, this is probably an underestimate of the true response rate, since the number of responsive genes for each TF was set to the minimum of the number in the TFKO and ZEV data sets. Individual rank response plots for all 11 other TFs present in all four data sets are shown in Figure S7.

**Figure 6.**
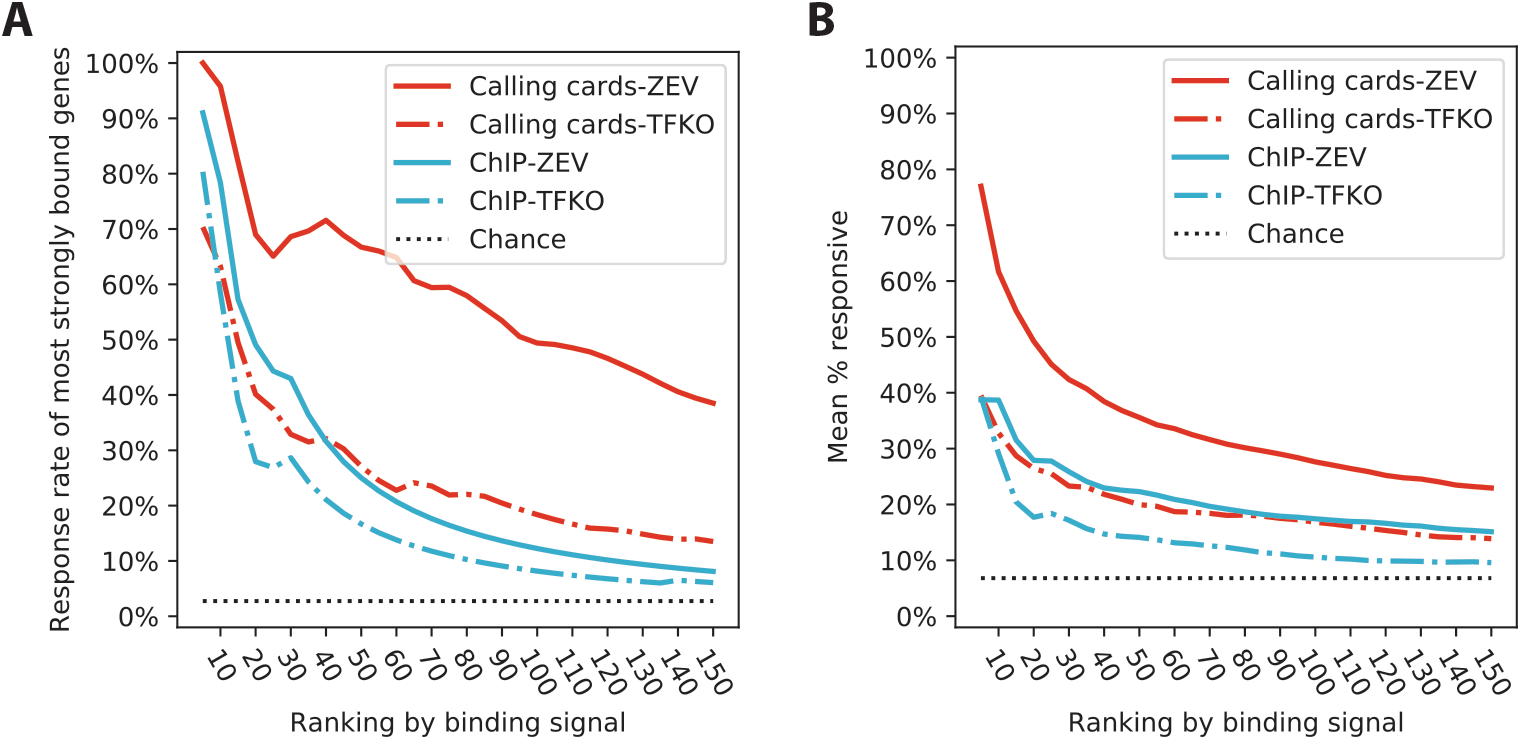
(A) The fraction of most strongly Leu3-bound genes that are responsive to Leu3 perturbation, as a function of the number of most-strongly bound genes considered. (B) Same as (A), with response rates averaged across the 12 TFs for which Harbison ChIP, calling cards, TFKO, and ZEV data were available.

## DISCUSSION

The fundamental question behind this investigation is how best to map the direct functional targets of transcription factors. We found that the established method of assaying the DNA binding locations of TFs, chromatin immunoprecipitation, does not by itself effectively identify the direct functional targets of a TF, because most of the genes whose cis-regulatory DNA is bound by a TF are not functionally regulated by that TF. We found this to be the case for two yeast ChIP datasets as well as 68 ENCODE ChIP-Seq experiments in human K562 cells and 88 ChIP-Seq experiments in human HEK293, consistent with previous reports based on different data sets. [15, 27, 28]

If the problem is that most bound genes are not responsive, a natural solution would be to focus on those that are. That is, to take the intersection of the genes a TF binds and the genes that respond to perturbation of the TF as its direct functional targets. However, we proved that this procedure does not effectively identify the direct functional targets when the sets of bound and responsive genes are much larger than their intersection. The reason is that, when there are many genes with non-functional binding sites and many genes that respond to the perturbation because they are indirect targets, it is expected that some genes will be indirect targets with non-functional binding sites in their cis-regulatory DNA. These are not direct functional targets, yet they inhabit and contaminate the intersection of bound and responsive genes. As a result, it is not safe to assume that genes that are both bound and responsive are responsive because they are bound.

We quantified this problem by setting minimal criteria for considering the genes that are bound and responsive to be likely direct functional targets. First, the intersection procedure must be able to achieve, in principle, 80% sensitivity with an expected false discovery rate of no more than 20%. Second, the intersection between the bound set and the responsive set must be greater than would be expected by chance, with a P-value of 0.01. We call a TF *acceptable* if it meets both those criteria. This designation does *not* guarantee that all or most of the TF’s bound and responsive genes are direct functional targets, i.e. that they are responsive because they are bound. In particular, the 80-20 criterion is a lower bound on the expected FDR, carrying no implications of any upper bound. Furthermore, it does not guarantee a *unique* relationship between the bound and responsive sets of an acceptable TF -- the bound set of one TF can be acceptable when compared to the responsive set of another TF, so long as the two sets show concordance beyond what would be expected by chance. *Acceptable* simply means that there is no obvious red flag to prevent us from supposing that a good number of the TF’s bound and responsive genes are direct functional targets. We found that, when combining ChIP data with steady-state perturbation-response data, the number of acceptable TFs was quite low. In both the yeast data and the human data, no more than 15% of the TFs assayed were acceptable. For the remainder, there is a clear red flag.

So far, we have assumed that any protein that is designated as a TF in the ENCODE database and has bound targets in ChIP-Seq and responsive targets in RNA-Seq is a TF. However, when we compared these to a recent, exhaustive, manually curated list of human TFs [29], we found that 20 TFKD targets and 11 CRISPRi targets were not on the list. One possible explanation is that these are sequence-specific DNA binding proteins that should have been on the Lambert list. A second possibility is that, although they do not bind DNA directly, they have ChIP-Seq peaks because they associate with proteins that do. A third possibility is that ChIP-Seq peaks do not necessarily reflect specific association with DNA, as suggested by a study in which green fluorescent protein (GFP) with a nuclear localization signal was found to generate thousands of robust ChIP peaks [30].

We identified four techniques that could increase the number of acceptable TFs substantially.

1. Measuring the transcriptional response a short time after inducing a perturbation by using a method such as ZEV. Overexpression by the ZEV system may also allow TFs with low activity in the experimental growth conditions to elicit a response from their target genes.
2. Using dual threshold optimization to set significance thresholds for binding and response data in a way that makes their intersection as significant as possible. This approach considers the two data types together, using each type to inform the threshold for the other, rather than considering each data type in isolation. Considering all the data should, logically, yield a better decision than only considering part of it, and we show that this approach does indeed yield more acceptable TFs.
3. Processing all the perturbation-response data together by using NetProphet 2.0, rather than considering the response to each perturbation in isolation from all the others.
4. Measuring TF binding location by using transposon calling cards rather than ChIP.

We are currently applying all these methods together to yeast and we expect the result to be a significantly expanded, high confidence map of the yeast TF network. As for mammalian cells, calling cards [31], dual threshold optimization, and NetProphet have all been shown to work. For TF activity perturbation, highly specific genome-targeting systems have been developed and tested with a variety activation and repression domains [32] and linked to small-molecule inducers [33, 34]. However, the prospects for obtaining ZEV-like perturbation and calling cards binding data on large numbers of mammalian TFs remain uncertain.

Other new technologies for measuring TF binding locations have shown great promise [35], but have not yet yielded a sufficiently large, systematic data set, with matched perturbation-response data, for comparison to ChIP and calling cards using the methods of this paper. One such technology is DAMID, in which a DNA-methyltransferase is tethered to a DNA-binding protein and changes in DNA methylation relative to a control are assayed to determine binding location [36–38]. Another is CUT&RUN, in which an endonuclease tethered to an antibody against a TF enters permeabilized nuclei and releases the DNA bound by the TF, which diffuses out of the cell and is recovered for sequencing [39–41]. A promising approach for measuring perturbation-response in mammalian cells is to transfect cells with a library of constructs encoding guide-RNAs that target a variety of TFs and then use single-cell RNA-Seq to identify the TF perturbed and measure the response. Variants of this general approach include Perturb-Seq [42, 43], CROP-Seq [44], and CRISP-Seq [45]. We expect that, as these technologies mature, they will be used to produce large systematic data sets that can be analyzed using the methods described here.

Even when we apply the best combination of analytic and experimental methods, a large fraction of the genes whose regulatory DNA is significantly bound by a TF binds do not respond to a perturbation of that TF. Such non-responsiveness could be caused by any of several mechanisms.

- Insufficient occupancy -- rank response plots (Fig. 6) indicate that the most strongly bound sites are much more likely to be functional than sites that are less strongly (but still significantly) bound.
- Saturation -- if a gene is already expressed at its maximum possible level and an activator of that gene is induced, no response will be seen. However, if other TFs were removed, lowering the expression level of the gene, it would respond to the induction. The same situation arises when a repressor of an unexpressed gene is induced or an activator of it is depleted.
- Inactivity -- the TF may bind DNA even when the TF is in an inactive, or partially active, state. However, the ability of ZEV induction of Gal4 to activate galactose genes even in the absence of galactose and presence of glucose shows that overexpression can elicit a response to TFs that are not normally active.
- Compensation -- the regulatory network as a whole may compensate for the change in TF activity in a way that damps the effect of the initial perturbation. Measuring responses shortly after the perturbation should reduce the prevalence of such compensation, but some mechanisms can compensate very quickly. A simple example would be two essentially equivalent TFs that can bind to the same sites, so that the effects of perturbing one TF are buffered by the other. This was shown to be a contributing factor in a comparison of the Harbison ChIP data to the TFKO data from Hu et al [15, 27]. Another example would be a TF that activates a protein that covalently inactivates the TF, such as a kinase or phosphatase.
- Override -- some regions of a genome may be shut down in a way that overrides the effects of TFs, even when the TFs can bind to the cis-regulatory DNA. For example, the transcribed region of a gene might be in inaccessible, tightly compacted DNA even though the cis-regulatory region remains somewhat accessible to TFs.
- Synergistic regulation -- some TFs that are bound to cis-regulatory DNA may be active only where there is a binding site for a cofactor nearby.

Regardless of the mechanism that renders a bound gene non-responsive, it remains the case that many binding sites are non-functional under the conditions tested, in the sense that the transcription rate of the associated gene is unaffected by the presence or absence of the TF. Currently, we do not know how much each of the factors listed above contributes to explaining why so many genes that are bound by a TF do not respond to a perturbation of that TF. For now, technical limitations of the available data sets may be a significant contributing factor. Once those have been mitigated by newer methods like transposon calling cards, we will be in a strong position to investigate the biological factors that explain the non-responsiveness of genes whose cis-regulatory DNA is bound by a TF. Determining the prevalence of each factor will bring the landscape of transcriptional regulation into much clearer focus.

## CONCLUSIONS

ChIP data on TF binding locations do not agree well with the set of genes that respond when the TF is genetically deleted (TFKO), confirming earlier findings. For most TFs, intersecting the bound and responsive genes is unlikely to be an accurate method of identifying direct functional targets. Agreement is improved by measuring binding using transposon calling cards and measuring response shortly after inducing the overexpression of a TF. When calling cards data become available for the entire set of yeast TFs, and ZEV induction-response experiments have been done in the same growth conditions, we will have a much clearer view of the network that regulates transcription in yeast.

## Supporting information

Supplementary File 1: Network

Supplementary File 2: Network

Supplementary File 3: Network

Supplementary File 4: Network

Supplementary FIle 5: Network

Supplementary File 6: Network

Supplementary FIle 7: Network

Supplementary FIle 8: Network

Supplementary File 9: TF lists

## METHODS

Detailed methods and data download links can be found in the online supplement.

## LIST OF ABBREVIATIONS

ChIP: chromatin immunoprecipitation
CRISPRi: CRISPR interference -- a method of repressing gene expression
DF: direct functional
DTO: dual threshold optimization -- a method of setting significance thresholds for binding and perturbation response-data targeting the same TF
TF: DNA-binding transcription factor
TFBS: TF binding site
TFKD: TF knockdown, encompassing siRNA and shRNA knockdowns
TFKO: TF knockout

## DECLARATIONS

Ethics approval and consent to participate: Not applicable. Consent for publication: Not applicable. Data availability: Data from the Harbison and Venters ChIP studies and Kemmeren TFKO studies are publicly available and can be obtained as described in refs. [14, 16, 23], respectively. ENCODE data can be obtained from the ENCODE web site: www.encodeproject.org. ZEV data are available at http://candid.research.calicolabs.com/. Calling Cards data are available as Supplementary File 10 in the peer reviewed, published paper.

The authors declare that they have no competing interests.

## FUNDING

MB was supported by NIH grants AI087794 and GM129126. YK was supported by matching funds for T32 HG000045 provided the McKelvey School of Engineering at Washington University. RM was supported by NIH grants GM123203, HG00975, and MH117070.

## AUTHORS’ CONTRIBUTIONS

MB conceived all computational analyses and drafted the ms. YK and NRP carried out the computational analyses, made figures, and drafted Methods. RSM conceived the ZEV experiments. BJW, GK, and RSM carried out ZEV experiments. PSR, XC, CS, and RM conceived, designed, and carried out the transposon calling cards studies.

## AUTHORS’ INFORMATION

YK is a computer science PhD student supervised by MB. NRP is a computer science master’s student supervised by MB. CS is a postdoctoral fellow supervised by RM. PSR is a postbaccalaureate fellow supervised by CS and RM. XC is a staff scientist. RM is the Goldfarb Professor of Genetics at Washington University School of Medicine. RSM, BJW, and GK are employees of Calico Life Sciences LLC.

## Notes

http://pin.research.calicolabs.com/

