## SupplementaryTextAndFigures for "Mapping Transcription Factor Networks By Comparing Tf Binding Locations To Tf Perturbation Responses"

### SUPPLEMENTARY FILES

1. Network edges from acceptable TFs to their bound and responsive genes after dual threshold optimization on Harbison ChIP and Kemmeren TFKO data.
2. Network edges from acceptable TFs to their bound and responsive genes after dual threshold optimization on Harbison ChIP and ZEV15 data.
3. Network edges from acceptable TFs to their bound and responsive genes after dual threshold optimization on Harbison ChIP and output from NetProphet run all Kemmeren expression data.
4. Network edges from acceptable TFs to their bound and responsive genes after dual threshold optimization on Harbison ChIP and output from NetProphet run ZEV 15 min, 45 min, and 90 min expression data.
5. Network edges from acceptable TFs to their bound and responsive genes after dual threshold optimization on calling cards and Kemmeren TFKO data.
6. Network edges from acceptable TFs to their bound and responsive genes after dual threshold optimization on calling cards and ZEV15 data.
7. Network edges from acceptable TFs to their bound and responsive genes after dual threshold optimization on calling cards and output from NetProphet run all Kemmeren expression data.
8. Network edges from acceptable TFs to their bound and responsive genes after dual threshold optimization on calling cards and output from NetProphet run ZEV 15 min, 45 min, and 90 min expression data.
9. TFs considered for individual networks: yeast TFs are proteins annotated with DNA binding domains; human K562 TFs are proteins ChIPped in major cell lines K562 and HepG2; human HEK293 TFs are zinc finger proteins ChIPped.
10. Calling Cards data (available in the peer reviewed, published paper).

### SUPPLEMENTARY FIGURE LEGENDS

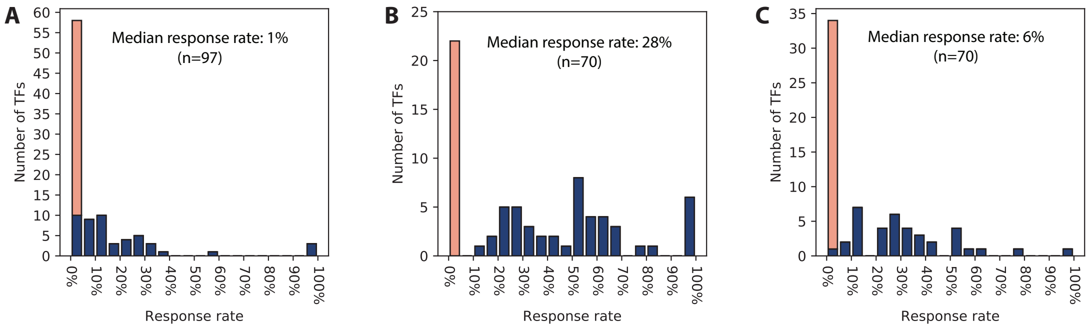


**Figure S1**. Same as Fig. 1 except: (A) Binding threshold is p<0.001 and response threshold is p<0.05 with fold change > 1.5. Total bound and responsive genes is 209. (B) Binding threshold is p<0.00001 and response threshold is p<0.05 with no minimum fold change. Total bound and responsive genes is 297. (C) Binding threshold is p<0.00001 and response threshold is p<0.05 with fold change > 1.5. Total bound and responsive genes is 119.

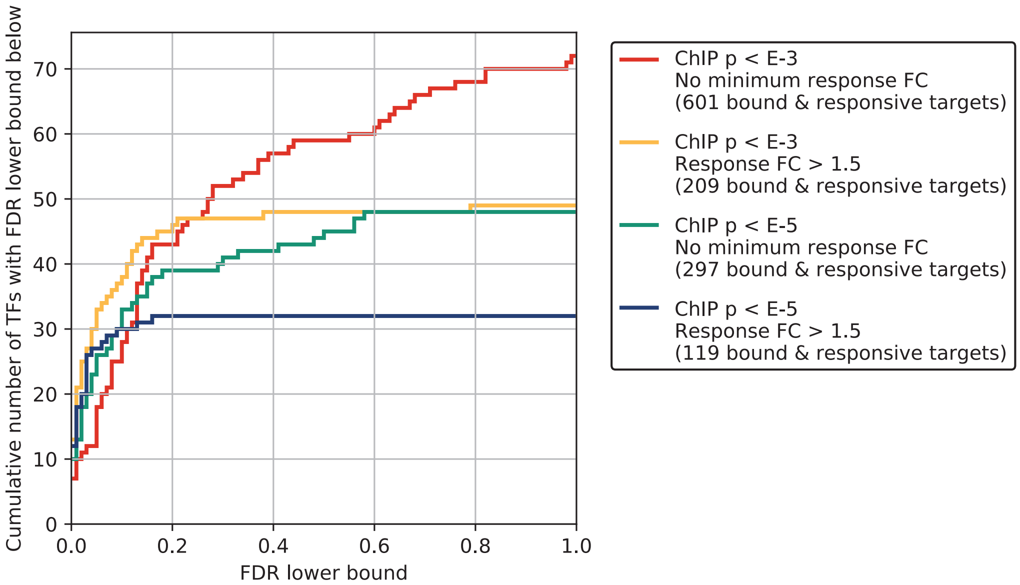


**Figure S2.** Cumulative number of TFs with expected FDR lower bound less than the number on the horizontal axis, assuming sensitivity of 80% (see Box 1). Red line: moderate binding and response thresholds; orange line: moderate binding threshold and tight response threshold; blue line: tight binding threshold and moderate response threshold (green), and tight binding and response thresholds.

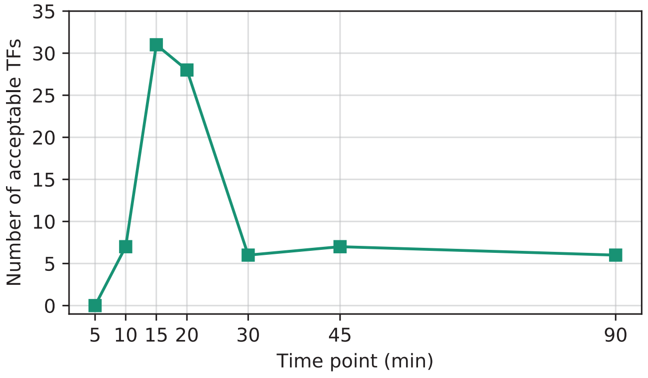


**Figure S3.** Numbers of acceptable TFs when comparing Harbison ChIP data to ZEV response data at various time points.

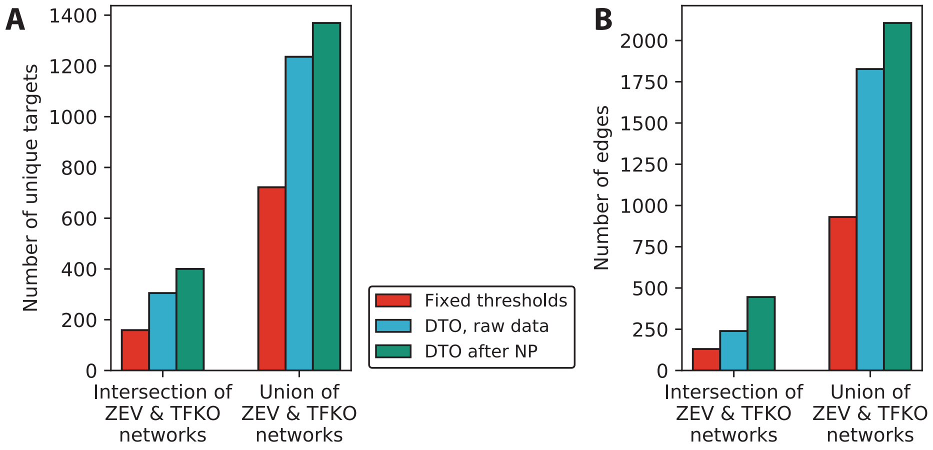


**Figure S4.** Comparison of TFKO and ZEV15 networks derived from fixed thresholds, dual threshold optimization (DTO) on raw gene expression, and DTO on gene expression data processed by NetProphet 2.0. The use of DTO on the raw expression data (blue bars) increases the size of the networks over intersection of bound and responsive sets determined by fixed thresholds (red bars). This is true whether you focus on the intersection of the TFKO and ZEV networks (left bar grouping) or the union (right bar grouping). Post processing expression data with NetProphet 2.0 (green bars) increases the further increases the size of the networks. The number off acceptable TFs is plotted in Fig. 3, unique target genes in Fig. S4A, and regulatory edges in Fig. S4B.

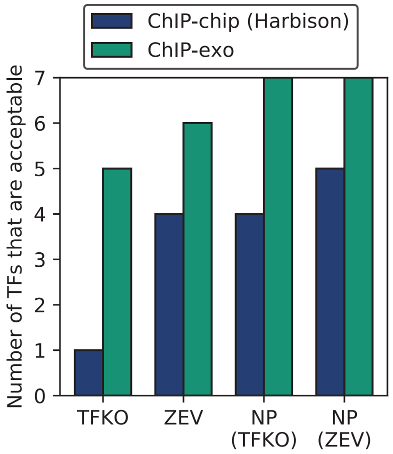


**Figure S5.** Among the 7 TFs for which we have data in Harbison ChIP, ChIP-exo, TFKO, and ZEV, the percentage that are acceptable. Regardless of the perturbation data set or processing by NetProphet 2.0, ChIP-exo in nitrogen-limited chemostats always yields more acceptable TFs than traditional ChIP-chip. NetProphet postprocessing always yields more acceptable TFs than raw differential expression.

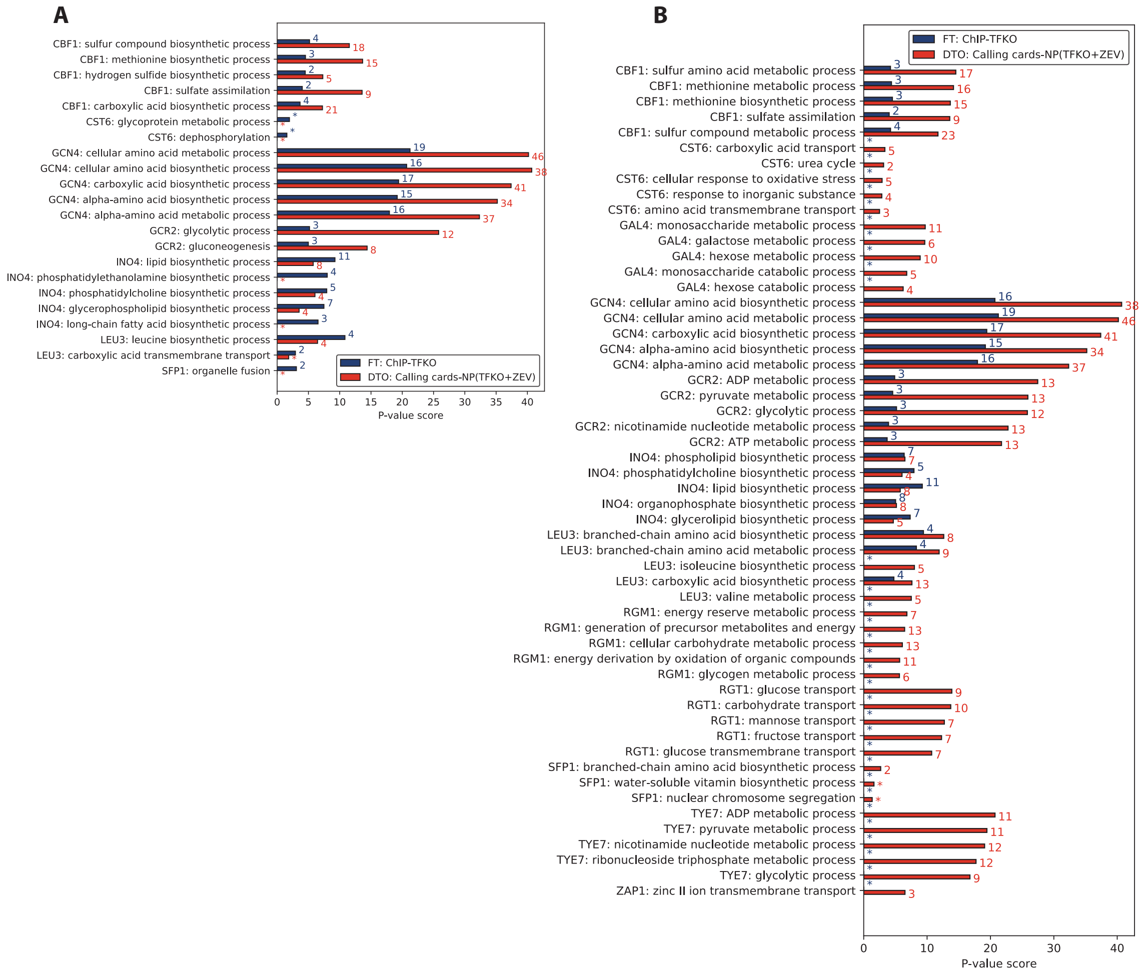


**Figure S6.** Same as Fig. 5C except: (A) For each TF that has any enriched GO term, the five most enriched terms (if available) when the targets of each TF are chosen by using fixed thresholds on Harbison ChIP and TFKO data. In the majority of cases, the most significant terms chosen by this method (blue bars) are even more significantly enriched when the targets are chosen by using dual threshold optimization on calling cards data and output from NetProphet 2.0 run on the TFKO and ZEV expression data (red bars). The numbers to the right of the bars indicate the number of genes with a given GO term among the targets of the TF. (B) For each TF that has any enriched GO term, the five most significantly enriched terms (if available) chosen by using dual threshold optimization on calling cards data and output from NetProphet 2.0 run on the TFKO and ZEV expression data. In many cases, terms with one or two fewer genes are more familiar than the terms with the most genes. For example, Gcr2 has 12 target genes annotated with “glycolytic process”, corresponding to its accepted function, but the most significant term is “ADP metabolic process”, which contains those 12 glycolytic genes plus one additional gene.

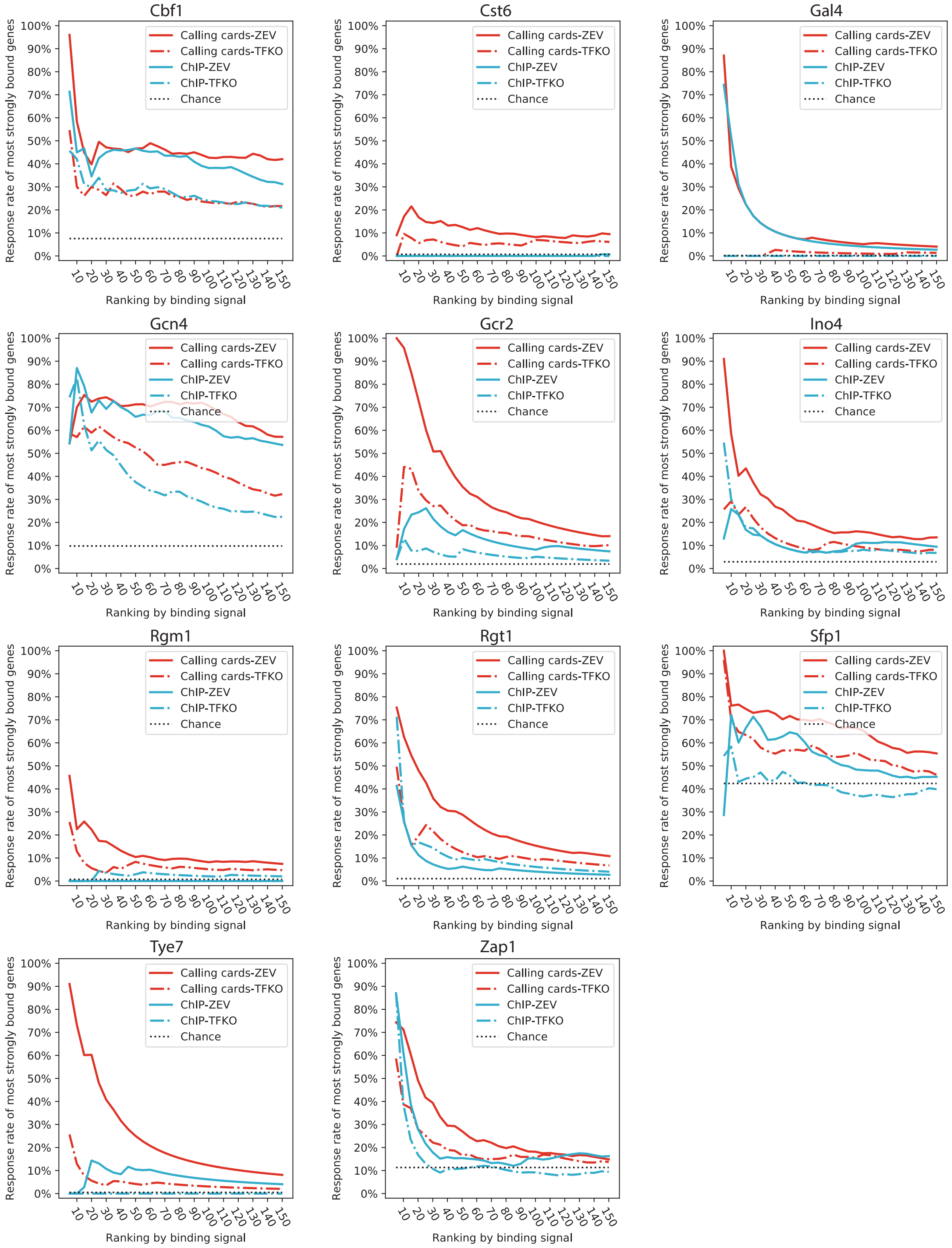


**Figure S7.** Same as Fig. 6A. The fraction of most strongly TF-bound genes that are responsive to the perturbation of that TF, as a function of the number of most-strongly bound genes considered. Shown are the other 11 TFs, in addition to Leu3, that had data available in Harbison ChIP, transposon calling cards, TFKO, and ZEV15.

### SUPPLEMENTARY METHODS

#### DATA PREPARATION

Yeast gene and TF definitions

For all yeast analyses, we considered the 5,887 genes labeled as "ORF verified" or "uncharacterized" in in the Saccharomyces Genome Database (SGD), discarding the 1,127 labeled as "dubious", "ncRNA", "rRNA", "snoRNA", "snRNA", or "tRNA". For TFs, we only counted those with evidence of direct DNA binding via a DNA binding domain, since they are the proteins that select the regulatory targets of regulators that bind indirectly. To identify these, we compared multiple lists, including those that had been ChIPed by Harbison et al., those that were over-expressed in the ZEV data, and those that had DNA binding specificity models in the CIS-BP database [1]. In cases of disagreement, we curated the list manually by consulting data in SGD, focusing primarily on domain analysis of the protein and on gene ontology categories assigned via high-throughput experiments such as protein-binding microarrays. In most cases the judgment is clear but there are some borderline cases that require a best guess. The complete list of 183 yeast proteins that we treated as TFs is provided as supplemental file 9.

Yeast ChIP-chip data sets

The Harbison ChIP-chip binding location data was published in [2]. We downloaded the p-values that represent the significance of TF binding within the intergenic regions from http://younglab.wi.mit.edu/regulatory_code/GWLD.html. The significantly bound targets were required to have $p < 0.001$ according to the authors' recommendation. TFs with no significantly bound target were eliminated from further analysis. The Venters ChIP-chip data were published in [3]. We downloaded the occupancy-level profiles for 200 transcription-related proteins from Table S4a in [3]. The occupancy level was the measure of log2 fold change of experimental signal over background signal within each regulatory region. The probes covered a distal region (260-320 bp upstream of ATG) and a proximal region (30-90 bp upstream of ATG). The downloaded occupancy level took the maximal level from either regulatory region. The authors used an FDR threshold of 5%, but we used 1% in order to make the data more comparable to those from the Harbison data set. For each TF, an FDR cutoff was calculated by searching for an occupancy level such that the ratio of number of targets in the mock IP control over the number in the experimental sample reaches the desired FDR. The "25&37C merged MockIP controls" file was obtained directly from the authors as it was unpublished. TFs with no significantly bound target were eliminated from further analysis.

(build: 19-Jan-2007)

Yeast ChIP-exo data sets

The ChIP-exo data for 12 TFs were compiled from four resources [4] [5] [6] [7]. We downloaded genomic coordinates of the ChIP peaks of Reb1, Gal4, Phd1 and Rap1 published in ref. [4] from https://ars.els-cdn.com/content/image/1-s2.0-S0092867411013511-mmc2.xls. We mapped the peaks to genes using the coordinates of each gene's promoter region (700 bp upstream to ATG) in reference genome S288C-R55, which was the closest last release prior to the date

cited in the paper, “(build: 19-Jan-2007)” (ref. [8] lists all releases). We then calculated the TF’s binding strength at each promoter as the sum of occupancy levels of all peaks assigned to that promoter. We downloaded peaks for Abf1 and Ume6 generated using a newer protocol, ChIP-exo 5.0, described in ref. [5], from GEO Series GSE110681. We also downloaded peaks for Cbf1 from GEO Series GSE93662 (see ref. [6]). The assignment of peak-promoter and binding strength at promoter were calculated as for the data from Rhee, et al., 2011, except that both peak and
promoter coordinates were based on gene annotation from reference genome S288C-R64. Lastly, we downloaded ChIP-exo data for Hap1, Ino4, Oaf1 and Pip2 from https://msystems.asm.org/content/msys/3/4/e00215-17/DC4/embed/inline-supplementary-material-4.xlsx (see ref. [7]). Each TF was assayed in four different environmental conditions, but we focused on the nitrogen limited chemostat data as that gave the best agreement with the expression data. The downloaded data set contains a score for each promoter that had at least one peak assigned to that promoter. The score represents the highest occupancy of those peaks assigned. Any promoter with no peak has a score of zero. We directly used the highest occupancy at each promoter as the binding strength, after removing any peak that was > 700 bp upstream from ATG.

Yeast transposon calling cards data

We used a published calling cards dataset on TFs Gal4 and Rgm1 [9], combined it with new data on Cbf1, Cst6, Eds1, Gcn4, Gcr1, Gcr2, Ino4, Leu3, Lys14, Rgt1, Sfp1, Tye7, Zap1. For ease of access, we submitted the combined data sets as Supplementary File 10. For each TF, all data from all replicates were combined. Transpositions within the promoters of yeast genes (700 bp upstream to ATG, reference genome S288C-R61) were used for calculating the significance of TF binding. For each promoter, a Poisson p-value was calculated by comparing experiment sample with no-TF control sample as described in [9]. To obtain a ranking by calling cards signal strength for dual threshold optimization, we used the normalized transposition count of the experimental samples minus that of the control samples to break ties when promoters had identical P-values.

Yeast TFKO data

The microarray expression data on gene knockout strains was published in ref [10]. The gene expression profiles of 1,484 single gene deletion strains and 3 wild type replicates were downloaded from http://deleteome.holstegelab.nl/data/downloads/deleteome_all_mutants_controls.txt. In addition, we downloaded the gene expression profiles after slow growth signature removed from http://deleteome.holstegelab.nl/data/downloads/deleteome_all_mutants_svd_transformed.txt. This transformed data set did not contain new p-values and analyzing it did not produce better results than the untransformed data, so we focused on the untransformed data.

Yeast ZEV induction data

The ZEV induction system was described in ref. [11]. The shrunken expression profiles were used as the quantitative responsiveness of target genes after TF induction. Specifically, the file called " Raw & processed gene expression data" was downloaded from http://candid.research.calicolabs.com/data and the column labeled `log2_shrunken_timecourses` was used. The responsive set contains all targets with non-zero expression levels. We systematically analyzed ZEV expression profiles measured at all time points (5, 10, 15, 20, 30, 45, and 90 minutes) after TF induction. To make different time points comparable, we only focused on 103 TFs that were commonly available in Harbison ChIP-chip dataset and each time point of the ZEV dataset. The maximal number of acceptable TFs was obtained at 15 min, so we chose to move forward with this time point for all subsequent analyses.

Human ChIP-seq data

Two human ChIP-seq data sets were analyzed in this work: ChIP-seq in K562 cell line published by ENCODE, and ChIP-seq in HEK293 cell line published in ref. [12]. All ENCODE data were downloaded from www.encodeproject.org as of January 21st, 2019. We focused on the data on K562 cells because it had by far the most TFs with both ChIP-seq and perturbation response data. We downloaded the “conservative” ChIP-seq peaks mapped to GRCh38 as called by the ENCODE pipeline, which uses the Irreproducible Discovery Rate (IDR) analysis of biological replicates with 2% IDR cutoff. Using the ENCODE definition of transcription factor, there was ChIP-seq data 261 TFs in K562. To quantify the significance of each TF-target binding interaction, we summed the log10 q-values of significant peaks that were within the regulatory regions of each gene (defined below). For the HEK293 cell line, we downloaded the combined summits for 131 ChIPped zinc finger proteins from GEO Series GSE76494 (see ref. [12] for details) The binding strength within the regulatory regions of a gene was the summed scores of all summits assigned to those regions. We tried two definitions of regulatory region: (1) a single long promoter extending from 10 Kb upstream of the 5’-most transcription start site (TSS) to 2Kb downstream (Ensembl Release 92), or (2) a core promoter extending from 500 bp upstream of the TSS to 500 bp downstream combined with the gene’s enhancers in the GeneHancer database [13] (V4.8). We used only the “double elite” enhancers, for which both the existence of the enhancer the gene-enhancer association are supported by at least two evidence sources. This double-elite list was obtained by emailing the authors of the paper. In order to properly use the ChIP summits in HEK293 whose coordinates were based on GRCh37, we used the LiftOver tool in UCSC genome browser to lift over the coordinates of regulatory regions from GRCh38 to GRCh37.

Human TFKD, CRISPRi and TF-induction data

We considered three human perturbation response data sets: TF knockdown (TFKD) in K562, CRISPRi in K562, and TF-induction in HEK293 [12]. The RNA-Seq expression profiles of wild-type controls, TFKDs and CRISPRi were downloaded from the ENCODE web site. Knockdowns using small-interfering RNA (siRNA) or small-hairpin RNA (shRNA) were combined in the data set we referred to as TFKD while the CRISPRi data were treated separately. For K562 cells, there were TFKD experiments targeting 261 different proteins and CRISPRi experiments targeting 96 different proteins. The expected counts were reported by RSEM program in the ENCODE RNA-seq processing pipeline using gene annotation from GENCODE V24 (GRCh38). Differentially expressed genes in each perturbed TF strain were processed by comparing the experimental replicate set to the control set using DESeq2 (V1.10.1). On the TF-induction set for HEK293, the processed RNA-seq expression profiles (after lowly-expressed gene removal and batch normalization) for 80 induced zinc finger proteins were downloaded from GEO Series GSE76495. Without control replicates, the expression levels of each profile were normalized to the medians of respective batch, as described in ref. [12].

NETPROPHET ANALYSIS

NetProphet 2.0 is a TF network inference algorithm that exploits gene expression data under genetic or environmental perturbation and genome sequences with annotations. The algorithm is described in detail in ref. [14]. Here, two yeast TF networks were mapped, one using the Kemmeren TFKO gene expression data and one using the ZEV data. Three human TF networks were mapped using the three perturbation response data sets.

Yeast NP networks

NetProphet 2.0 requires gene expression data in the form of a gene expression matrix and differential expression matrix. The Kemmeren gene expression matrix was represented as the log2 fold-change (logFC) values of strains with gene deletions over wild-type strains. The ZEV gene expression matrix was represented as the logFC values of the levels measured at a certain time point after the TF induction relative to time 0. For the differential expression (DE) module of the algorithm, we used Kemmeren samples in which a TF-encoding gene was knocked out, not those in which some other type of gene was knocked out, and ZEV samples from 15 min after TF induction. For the co-expression module of the algorithm, we used the complete set of 1,484 Kemmeren expression profiles from strains lacking one gene (not necessarily encoding a TF) or 590 ZEV expression profiles from 15 minutes, 45 minutes, or 90 minutes post-induction. The other two inputs were DNA sequences of yeast promoters and amino acid sequences of TFs' DNA binding domains (DBDs), as described in ref. [14]. PWM models of TFs' binding specificity were not used. Each output network is an adjacency matrix, where the rows represent TFs, the columns represent genes, and the entries are NetProphet scores representing the aggregate strength of evidence that the gene is a direct functional target of the TF.

Human NP networks

For K562 data, we calculated differential expression (DE) P-values for TFKD and CRISPRi independently using DESeq2. The DE matrix input to NetProphet 2.0 contained the -log P-values, with a negative sign for apparent repression (the knockdown of the TF makes the target gene go up). For HEK293 data, we directly used the logFC values because there were no replicates, hence no P-values, for most TFs. The co-expression matrix contained the logFC of individual mutant strain replicates over the median expression level of control replicates (K562) or the median expression level of the gene across all perturbations (HEK293). There were 765 expression profiles (including replicates) for K562 TFKD, 252 for K562 CRISPRi, and 107 for HEK293 TF inductions. We obtained DNA sequences of the regulatory regions based on their coordinates in GRCh38 using our definition (2) of regulatory regions. We concatenated the enhancers and promoter of each gene into a single sequence for the purpose of motif inference in NetProphet 2.0. Each pair of concatenated regions was separated 50 N's to ensure that no inferred motif instances crossed between one enhancer and another. We also queried the CIS-BP [1] database for the amino acid sequences of human TFs' DBDs. The detail of DBD preprocessing is described in [14]. The TFKD and CRISPRi networks had 392 TFs each, which were ENCODE TFs being ChIPped in either of the major cell lines K562 or HepG2. The TF-induction network had 103 TFs, which were the zinc finger proteins being ChIPped in HEK293.

### ACCEPTABLE TFS

For each pair of binding and expression data on a given TF, the positive gene sets (bound genes or responsive genes) were compared. The TF was deemed acceptable if they met two criteria. (1) The lower bound on the expected FDR had to be less than or equal to 20% when the sensitivity was fixed at 80%, as calculated by using the formula derived in Box 1. (2) The P-value for the significance of the overlap between the bound and responsive gene sets had to be <= 0.01. For fixed threshold analysis, the P-value was calculated using the hypergeometric null distribution. For dual threshold analysis, it was calculated using the randomization-based null distribution, not the nominal P-value (see description of dual threshold optimization).

### DUAL THRESHOLD OPTIMIZATION

#### Software availability

Software implementing dual threshold optimization and instructions can be found at https://github.com/patelni14/dual_threshold_optimization.

#### DTO algorithm

For each TF, dual threshold optimization (DTO) uses one binding location data set and one gene expression data set. The genes in each data set are ranked by the strength of their binding or expression signal. By default, the signal strength is the negative log P-value, but different experiments and methods may require different calculations of signal strength (see sections below). For each data set, DTO chooses a threshold on the ranks such that genes ranking above the threshold are considered positives for binding or response (see Fig. 3C). A series of rank-threshold combinations (places where the gray lines cross in Fig. 3C) are used to generate positive subsets of genes in each dataset. The series of thresholds for each data set, $T_1, T_2, \ldots$, were generated using the recurrence:

$$T_1 = 1$$
$$T_n = Floor(T_{n-1} * 1.01 + 1)$$

This formula produces a fine spacing among smaller subsets that becomes coarser as the subsets grow. If a threshold would split a group of genes that all have the same score that threshold is skipped. For each pair of subsets, a hypergeometric p-value was computed using a hypergeometric survival function (Scipy's hypergeom.sf) with the following parameters:

k = # of genes in the intersection of the subsets - 1
M = # of genes in the universe of assayed genes
n = # of genes in the expression subset

N = # of genes in the bound subset

This hypergeometric P-value is the probability of an intersection as large as, or larger than, the observed intersection, when choosing random subsets of genes, with the number of genes in each random subset equal to the number of genes in the positives defined by the rank threshold pair. We refer to this as the *nominal P-value* because it is only used for selecting the best pair of thresholds, not for determining whether the resulting overlap is significantly larger than would be expected by running DTO on random rankings. DTO returns the threshold combination that minimizes the nominal P-value.

Randomization-based P-values for overlaps identified by DTO

To produce a null distribution for testing the significance of the overlap chosen by DTO, a randomization procedure was used. A new set of data was generated using random assignment of signal strength scores to genes and then DTO was run. The best nominal P-value for each randomized data set was used to calculate a null distribution of nominal P-values. This was done by running the randomized DTO procedure 1000 times for each TF in each analysis and the distribution of nominal (hypergeometric) p-values enabled us to determine a $P<0.01$ significance threshold on the nominal P-value that is specific to that TF. When the nominal P-value of the rank pair chosen by DTO using the true data was below the threshold defined by the randomizations, the overlap was considered significant.

Application of DTO to yeast data

In the analysis of yeast data, the universe was defined as the set of all genes assayed in either of the two datasets being compared. For data sets that do not have P-values, the signal strength is the log fold change (ZEV) or score (NetProphet). For Calling cards, where many P-values were identical, ties were broken by the difference between the number of insertions in the experimental sample and the number of insertions in the control sample. The number of insertions was normalized to the total insertion count in each sample.

Occasionally, DTO can produce implausible results, such as concluding that all genes are responsive to a perturbation. To prevent this, we set very relaxed limits on the bound or responsive genes in certain data sets. For Harbison ChIP, we required $P<0.1$. For NetProphet output we required that the score of the TF-target relationship be among the top 150,000 scores. These were sufficient to eliminate any anomalous results; no constraints on the TFKO or ZEV data were required.

Application of DTO to human ENCODE data

In the analysis of human data on K562 cells, the universe was defined as the set of all genes detected in the gene expression dataset. Response signal strength for DTO was the absolute value of the log fold change, relative to non-perturbed control samples. DTO was limited to choosing bound or responsive genes with $P<=0.1$. For NetProphet output we required that the score of the TF-target relationship be among the top 500,000 scores.

Application of DTO to human HEK293 data

In the analysis of human data on HEK293 cells, the universe was defined as the set of all genes detected in the gene expression dataset. Response signal strength for DTO was the absolute value of the log fold change, relative to non-perturbed control samples. No replicates or P-values were available for most TFs. For both raw perturbation-response data and NetProphet scores, we required that the score of the TF-target relationship be among the top 300,000 scores.

### COMPARISONS AMONG BINDING DATA SETS

#### Harbison ChIP-chip compared to Venters ChIP-chip

The TFs used in the comparison shown in Figure 5A are: ASH1, BAS1, CHA4, FKH1, FKH2, GCN4, GLN3, INO4, LEU3, MSN2, PHO2, RFX1, RPH1, RSF2, SFP1, SKN7, STE12, STP1, SWI5, UGA3, WTM1, WTM2, XBP1, YAP5, ZAP1

#### Harbison ChIP-chip compared to ChIP-exo

The TFs used in the comparison shown in Figure S5 are: CBF1, GAL4, HAP1, INO4, OAF1, PHD1, PIP2

#### Harbison ChIP-chip compared to transposon calling cards

The TFs used in the comparison shown in Figure 5B are: CBF1, CST6, GAL4, GCN4, GCR2, INO4, LEU3, RGM1, RGT1, SFP1, TYE7, ZAP1

### RANK RESPONSE PLOTS

To create the lines in the rank response plots such Fig. 6A, we first determined the minimum of the number of responsive genes in the TFKO and the ZEV data -- call it n. A gene was considered responsive in the TFKO data if it had adjusted $P<0.05$ and in the ZEV data if it had shrunken absolute log fold change > 0. We then labeled the top n most strongly responsive genes in the TFKO and ZEV data as responsive for purposes of this plot (see "DTO algorithm" above for definitions of signal strength). This equalized the number of ZEV-responsive and TFKO-responsive genes for each TF. We then sorted genes by the strength of their binding signal for the TF in question. Next, we considered the top 1, 2, 3, 4, etc. most strongly bound genes. For each such group, we calculated and plated the fraction of genes that were responsive. For the mean rank-response plot (Fig. 6B) we simply averaged the response rates across the 12 TFs.

### GO ENRICHMENT ANALYSIS

Gene ontology (GO) enrichments for each TF were analyzed using two networks mapped using different methods: (1) fixed threshold on Harbison ChIP data and TFKO data; (2) DTO on calling cards data and output from NetProphet 2.0 run on the TFKO and ZEV expression data. For each TF, it target set in network 2 was the union of the output of DTO applied to NetProphet scores from ZEV expression data and DTO applied to NetProphet scores for TFKO expression data. The mapping of GO term to gene for *Saccharomyces cerevisiae* was queried using R Bioconductor library org.Sc.sgd.db (V3.5.0). Any GO terms annotated with less than 3 or greater than 300 genes were eliminated. Using the target genes of a TF identified from network (1) or

(2), the GO enrichment in biological process was analyzed using hypergeometric test implemented in R Bioconductor library GOstats (V2.44.0). The output p-values were used for ranking the enriched terms, from most significant to the least significant. When plotting the top GO term shown in Figure 5C, for each TF, we combined all terms from both networks into a single rank list. If multiple terms were enriched by the same set of targets, only the children term (the most specific term) was retained using the GO hierarchical structure, i.e. those other ancestral (redundant) terms were removed from the rank list. Subsequently, the top GO term was chosen for the corresponding TF. When plotting the top GO terms shown in Figure S6, we used the GO terms enriched in one network and chose the top five (if available). The same redundancy removal strategy was applied.
